# Iterative SCRaMbLE for Engineering Synthetic Genome Modules and Chromosomes

**DOI:** 10.1101/2024.12.06.627136

**Authors:** Xinyu Lu, Klaudia Ciurkot, Glen-Oliver F. Gowers, William M Shaw, Tom Ellis

## Abstract

Synthetic biology offers the possibility of synthetic genomes with customised gene content and modular organisation. In eukaryotes, building whole custom genomes is still many years away, but work in *Saccharomyces cerevisiae* yeast is closing-in on the first synthetic eukaryotic genome with genome-wide design changes. A key design change throughout the synthetic yeast genome is the introduction of LoxPsym site sequences. These enable inducible genomic rearrangements *in vivo* via expression of Cre recombinase via SCRaMbLE (Synthetic Chromosome Recombination and Modification by LoxPsym-mediated Evolution). When paired with selection, SCRaMbLE can quickly generate strains with phenotype improvements by diversifying gene arrangement and content in LoxPsym-containing regions. Here, we demonstrate how iterative cycles of SCRaMbLE can be used to reorganise synthetic genome modules and synthetic chromosomes for improved functional performance under selection. To achieve this, we developed SCOUT (<u>S</u>CRaMbLE <u>C</u>ontinuous <u>O</u>utput and <u>U</u>niversal <u>T</u>racker), a reporter system that allows SCRaMbLEd cells to be sorted into a high diversity pool. When coupled with long-read sequencing, SCOUT enables high-throughput mapping of genotype abundance and correlation of gene content and arrangement with growth-related phenotypes. Iterative SCRaMbLE was applied here to yeast strains with a full synthetic chromosome, and to strains with synthetic genome modules encoding the gene set for histidine biosynthesis. Five synthetic designs for *HIS* modules were constructed and tested, and we investigated how SCRaMbLE reorganised the poorest performing design to give improved growth under selection. The results of iterative SCRaMbLE serve as a quick route to identify genome module designs with optimised function in a selected condition and offer a powerful tool to generate datasets that can inform the design of modular genomes in the future.

## Introduction

Over the last 15 years the field of synthetic genomics has advanced from the initial full synthesis and assembly of genomes to genome minimisation and genome-wide recoding projects in bacteria^1–3^. In eukaryotes, the Synthetic Yeast Genome (Sc2.0) project is nearing completion, with almost all of the 16 synthetic chromosomes completed individually, and many now merged into a single *Saccharomyces cerevisiae* strain^4–6^. But so far, the ‘synthetic-ness’ of these synthetic genomes has been somewhat limited, with the gene organisation of the assembled chromosomes and genomes always following that of the native version. As we push towards ever-more synthetic genomes, where the genes on chromosomes are organised into functional modules built-to-design from DNA parts^7^, we need to develop new methods that allow us to explore the ideal gene layout for a set of genes and to better understand how gene arrangement effects fitness and gene expression.

Recently we described the development of synthetic genome modules for yeast^8^. Going beyond the work of the Sc2.0 project, these modules are synthetic regions of chromosomes built to contain a set of genes that together encode a common function, with the native copies of those genes removed from the rest of the genome. When the genes in such modules relocate along with their flanking regulatory elements (promoters and terminators), they are known as ‘defragmented’ modules. But toolkits for modular DNA part assembly in yeast^9–11^ allow us to go further and make ‘refactored’ modules where the native elements like promoters are replaced with well-characterised synthetic counterparts, allowing us to test our understanding of how gene expression control contributes to the function of the module. Indeed, refactoring of endogenous genes is a well-established route to understanding regulation and has been achieved in regions of phage, bacterial and yeast genomes^12–16^.

So far the few studies that have tried defragmenting genes in genomes or refactoring them have not explored how arrangement of the genes effects their expression and function. In yeast, the Synthetic Chromosomal Rearrangement and Modifications by LoxPsym-mediated Evolution (SCRaMbLE) system, developed as part of the Sc2.0 project, offers an exciting tool for this this work. SCRaMbLE enables inducible genome rearrangement by Cre-mediated recombination between loxPsym sites, generating gene deletions, inversions, duplications^17–19^. The resulting changes in gene copy number, order and orientation in DNA constructs have been shown to drive transcriptional changes, which *in vitro* has been demonstrated to fine-tune gene expression to optimise function of a heterologous pathway^20^. However, due to the randomness of the diversity generated by SCRaMbLE, achieving maximal phenotypical improvements in a single round of SCRaMbLE is unlikely and often many rounds are needed. A method known as multiplex SCRaMbLE iterative cycle (MuSIC) was developed to address this, continuously generating genome diversification by SCRaMbLE cycles with screening for the most desired phenotypes done in a stepwise manner^21,22^. We anticipate that iterative rounds of SCRaMbLE (as used in MuSIC) while suitable for finding optimal gene arrangements, will plateau at solutions representing local maxima in the design space. Determining this is challenging, as resolving rearranged genotypes *en masse* is not straightforward with widely used short read DNA sequencing methods.

Most post-SCRaMbLE screening methods are low-throughput and rely on single colony analysis to confirm phenotypes along with genome sequencing to determine the rearranged genotypes and decipher their influence^22–25^. For iterative SCRaMbLE methods that involve extensive screening across multiple cycles, a high throughput method to map all genotypes of a post-SCRaMbLE population and quantify each genotype’s relative fitness is lacking. A further complication is that a significant percent of cells in a population of yeast induced to SCRaMbLE do not actually undergo any Cre-mediated rearrangements. These non-recombined yeasts can complicate iterative approaches and use up sequencing and screening capacity. To efficiently isolate SCRaMbLEd cells, a reporter system known as ReSCuES was previously developed to use auxotrophic selection to select against un-SCRaMbLEd cells^25^. ReSCuES flips a dual auxotrophic selectable marker cassette to toggle between two selections, enabling efficient screening of post-SCRaMbLE colonies by implementing targeted selection^25^. However, using this reporter system restricts selectable marker availability in yeast engineering and risks losing positive SCRaMbLE events as the marker is reversible with every Cre rearrangement. To free-up marker usage and maximise the capturing of positive SCRaMbLE events, developing an alternative selection system is necessary.

Here working in yeast, we explored the idea that iterative SCRaMbLE could be used as a tool for improving the design of refactored synthetic modules and genomes. To test this out, we first constructed synthetic genome modules encoding histidine biosynthesis and examined whether moving these genes into new synthetic arrangements led to growth and function defects. We then showed that SCRaMbLE could be used to rescue a defective *HIS* module by rearranging genes within this module, with this identifying optimal configurations under specific growth conditions. To facilitate post-SCRaMbLE screening in this work, we developed SCOUT (<u>S</u>CRaMbLE <u>C</u>ontinuous <u>O</u>utput and <u>U</u>niversal <u>T</u>racker) to allow us to efficiently isolate cells using FACS that are expected to have undergone SCRaMbLE. SCOUT was then combined with POLAR-seq^26^, to enable us to sequence a sorted pool of cells and resolve the relative fitness of each rearranged genotype by correlating its abundance with phenotype improvements. We then characterised the accumulation of rearrangements during successive rounds of SCRaMbLE, both within a synthetic genome module and across a synthetic chromosome, demonstrating in both cases how rearrangements improve phenotypes but quickly reach a plateau. Our study gives new insights into how gene rearrangements in synthetic yeast can improve selected functions and provides new tools for others to use SCRaMbLE to quickly achieve improved phenotypes from combinatorial libraries of rearranged genotypes.

## Results

### Design and construction of the synthetic *HIS* modules

To explore how gene rearrangement can impair or optimise gene expression in synthetic genomes, we first required a testbed system where such changes can be easily linked to a phenotype. In recent work we demonstrated the concept of synthetic genome modules by creating yeast strains with *TRP* modules that co-locate the genes encoding the tryptophan biosynthesis pathway into a synthetic cluster containing loxPsym sites between the genes^8^. Here, we applied our synthetic genome module approach to the genes encoding histidine biosynthesis in yeast. Seven *HIS* genes, namely *HIS1* to *HIS7*, catalyse the 10 reaction steps of the histidine biosynthesis from phosphoribosyl pyrophosphate (PRPP) to L-histidine (**Figure 1A**). These genes were relocated to synthetic modules in the *URA3* locus of the yeast genome following one of two approaches; either relocating genes with their native flanking regulatory elements - an approach we called *‘defragmentation*’, or by relocating only the protein coding sequences (CDS) of the genes and rebuilding them into a gene cluster with commonly-used modular promoter and terminator parts - an approach we call *‘refactoring’* (**Figure 1B**).

**Figure 1.**
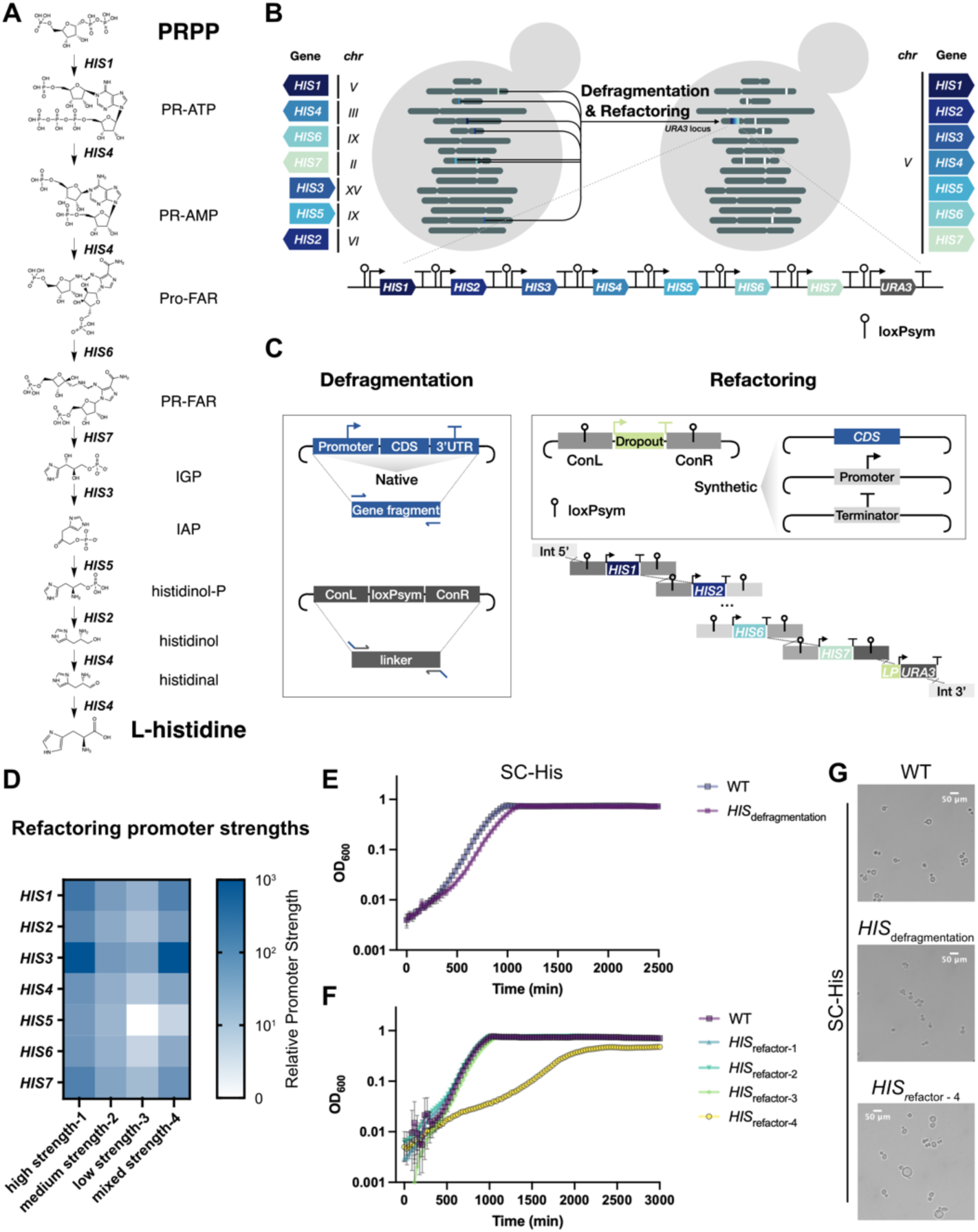
Design and construction of histidine genome modules. (A) Metabolic pathway of L-histidine biosynthesis from PRPP in *S. cerevisiae*. (B) Schematic overview of the relocation of *HIS* genes into a synthetic module by defragmentation and refactoring. Seven *HIS* genes catalysing 10 steps of reactions in the histidine biosynthesis pathway, namely *HIS1* to *HIS7*, including their native regulatory sequences, were partially removed from their native genomic loci and relocated as a module at the *URA3* locus. Genes are labelled in distinct colours. White squares labelled on chromosomes represent deletion of seven *HIS* genes from their native loci and replacement with a 23 bp ‘landing pad’ which contains a unique CRISPR/Cas9 targeting sequence. (C) Schematic process of the defragmented and refactored *HIS* module assembly. For defragmentation, gene fragments containing the native promoter (1 kb region upstream of the gene), CDS and 3’UTR (500 bp region downstream of the gene) of each *HIS* gene were amplified from the BY4741 genomic DNA and then constructed into entry-level plasmids by Gibson assembly. The linker plasmids were constructed by inserting a loxPsym sequence into the synthetic connectors from the yeast Moclo Toolkit (YTK)^10^. For refactoring, the CDS of each *HIS* gene was firstly constructed into an entry-level plasmid as a synthetic ‘part’ that is compatible for YTK assembly. Next, the YTK promoter, terminator along with the CDS of *HIS* genes were constructed as gene cassettes into the vectors containing linkers embedded with a 34 bp loxPsym sequence. Gene fragments and linkers for defragmentation, and gene cassettes with their linkers for refactoring, were linearised from the plasmids and then integrated into the *URA3* locus as a synthetic module by yeast homologous directed repair (HDR)-based assembly, respectively. (D) Heatmap representing the range of constitutive promoter strengths. Each promoter was assigned for a score from 0-1000 based on the data characterised in YTK^10^. Different combinations of promoters were selected for *HIS* gene expression cassettes. (E) Growth curves of the WT control strain and the strain harbouring the defragmented *HIS* module in SC-His, n=4. (F) Growth curves of the WT control strain and the strain harbouring the refactored *HIS* modules in SC-His, n=3. (G) Microscopy images of the WT control strain and strains harbouring the defragmented and refactored synthetic *HIS* module (*HIS*refactor-4) from the overnight culture in SC-His.

To relocate the *HIS* genes, we began with a BY4741 strain with the *HIS3* gene partially deleted^27^ and used three iterative rounds of CRISPR-mediated deletion to remove the coding sequences of the other six *HIS* genes and their immediate flanking regulatory sequences, being careful to avoid deleting any sequences that might act as regulatory elements of the neighbouring genes (**Figure S1**). Each deleted gene was replaced with an individual 23 bp ‘landing pad’^14^ designed so that it can be easily targeted with CRISPR/Cas9 for future gene restoration or further locus deletion. Successful gene deletion was confirmed by junction PCR from the genomic DNA of transformant colonies **(Figure S2A and S2B**).

To generate a strain with a defragmented *HIS* module, we first constructed the *HIS* genes with their native regulatory sequences as individual gene cassette plasmids (**Figure 1C**). These gene cassettes were designed to include around 1 kb upstream and 500 bp downstream sequence for each gene in order to preserve its native promoter and terminator. Synthetic linkers (∼200 bp) to connect the 7 gene cassettes into a module were also made by adapting YTK assembly connectors^10^ and designing these to all embed the 34 bp loxPsym sequence^5,17^. The seven *HIS* gene cassettes, eight synthetic linkers and a *URA3* selectable marker, were linearised from assembled plasmids and co-transformed into the yeast strain yXL010 with all *HIS* genes deleted. During the transformation all DNA parts were linked together in yeast by homology-dependent recombination, integrating as a 20 kb synthetic module at the *URA3* locus (**Figure S2C**). Junction PCR on genomic DNA of the transformants confirmed yeast colonies with the correct defragmented *HIS* module assembled into the desired locus (**Figure S2C-S2E**). One of these clones (yXL052) was selected for further examination.

In parallel to making the defragmented *HIS* module, we also built yeast strains with refactored *HIS* module. To do this we selected constitutive promoter and terminator parts used in the YTK system^10^ and assembled them with *HIS* gene CDS parts to replace their native promoters and 3’ UTRs (**Figure 1C**). For these modules we used a variety of promoters, classifying YTK promoters as either high, medium or low strength. We also ensured that we did not repeat any YTK parts within any module designs in order to avoid generating repetitive DNA that may trigger homologous recombination. We constructed four refactored module designs, with the first three being modules with all strong, all medium and all weak promoters (**Figure 1D**). In each case we chose which promoters from each classification should be matched to each of the 7 *HIS* genes by analysing native *HIS* gene transcription levels from previous RNA-seq assays^28^ and isoform profiling experiments^29^, This meant that the gene associated with the most transcription would be assigned with the strongest promoter from the classification (e.g. the strongest ‘weak promoter’) and the least transcription would use the weakest promoter.

Finally, to assess the pathway’s robustness to a broader range of gene expression variations we made a fourth ‘mixed strength’ combination where promoters for each *HIS* gene were selected from each of the 3 defined groups, with the strongest promoter assigned for *HIS3*, and one of the weakest promoters assigned for *HIS5* (**Figure 1D**). As with the defragmented module, these refactored modules were assembled from linear DNA parts in one go by using yeast assembly during transformation to link the DNA together by homology-dependent recombination and insertion into the *URA3* locus of the *HIS* deletion strain (**Figure 1C**). A CRISPR-targetable ‘landing pad’ was included in the module design to enable future module expansion or further editing at this genomic locus. Successful module assemblies were all confirmed by junction PCR from the genomic DNA of transformant colonies (**Figure S3**).

### Refactoring with imbalanced *HIS* gene expression causes a functional defect

We next assessed the functionality of the five synthetic *HIS* modules and fitness of the strains by comparing their growth to a wildtype (WT) control strain (yXL014) rich media (YPD) and synthetic complete media with (SC) or without histidine (SC-His). In these three tested conditions none of the strains except the one harbouring the mixed promoter strength refactored module, known as *HIS*_refactor-4_, exhibited any significant growth defects (**Figure 1E, F, Figure S4**). The *HIS*_refactor-4_ strain showed significantly enlarged budding cells and a slow growing phenotype in SC-His (**Figure 1F, G**). This same strain, however, exhibited a normal growth rate and cell size when grown in media with histidine present (**Figure S4B, D**). This indicates that the mixed promoter strength module design is specifically sub-optimal for histidine biosynthesis but is unlikely to be affecting cell health in any other way, despite the relocation of the genes.

By matching the promoter choice for each *HIS* gene in *HIS*_refactor-4_ to the corresponding enzymatic reactions of the pathway, we deduced that the slow growing phenotype in SC-His medium might be due to having a very weak promoter (*pRAD27*) for *HIS5* and a very strong promoter (*pTDH3*) for *HIS3.* This creates a possible 150-fold difference in enzyme expression between these two genes that theoretically should lead to the accumulation of the intermediate metabolite IAP during pathway function and a large reduction in the flux into the downstream reactions that produce the histidine needed in SC-His conditions. Given that all our synthetic *HIS* modules contained loxPsym sites between the genes, we hypothesised that this poor design for *HIS*_refactor-4_ could be automatically resolved *in vivo* by the SCRaMbLE system which can induce gene deletions, inversions, and duplications, through Cre-mediated recombination (**Figure 2A**). Thus, this strain (known as yXL219) offers a testbed for using SCRaMbLE as a method to rapidly change gene expression levels and in doing so identify module redesigns that will have improved performance.

**Figure 2.**
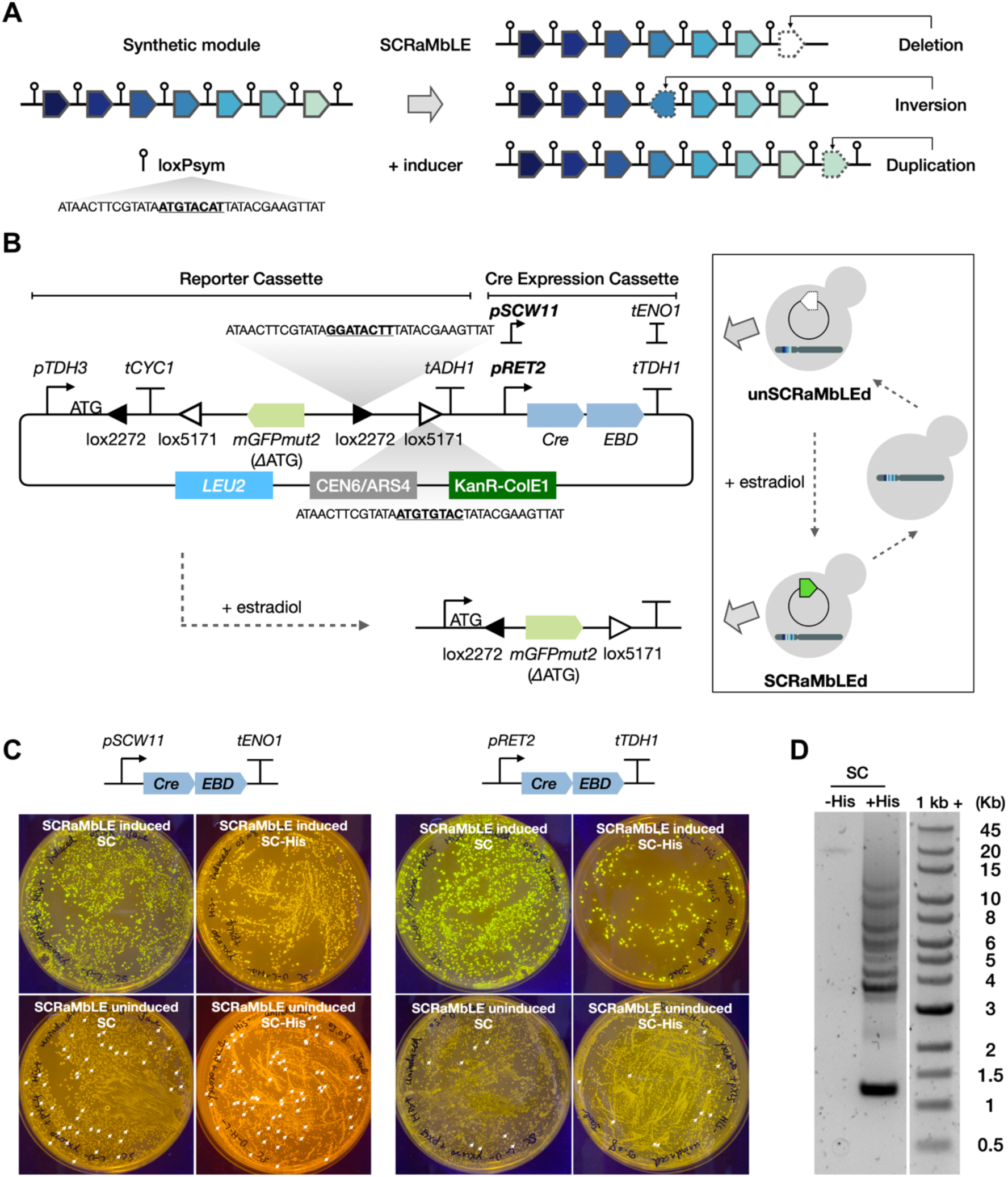
SCOUT for SCRaMbLE screening. (A) Schematic of SCRaMbLE inducing rearrangements, such as gene deletion, inversion and duplication between loxPsym sites in synthetic DNA. (B) Schematic of the SCOUT reporter design and its application in iterative SCRaMbLE. The SCOUT cassette is an antisense-orientated *mGFPmut2* gene lacking ATG start codon flanked by two pairs of loxP variant sites, loxP2272 and loxP5171. The reporter also encodes a CreEBD expression cassette, where Cre recombinase is fused to the oestrogen binding domain (EBD) so that its function can be induced by β-estradiol. Cre-mediated inversion and excision occur at each pair of loxP variant sites respectively, resulting in the stable inversion of the *mGFPmut2* gene and deletion of one of each pair of lox sites. The flipped *mGFPmut2* orientation turns on GFP fluorescence. The reporter plasmid can be removed from SCRaMbLEd strains and reintroduced for a new cycle of SCRaMbLE. (C) Assessment of the SCRaMbLE reporter performance. SCRaMbLE was induced by adding 1 *μ*M β-estradiol to a strain with a defragmented *HIS* module. After 4 hours cells were washed twice and plated on SC and SC-His plates respectively at a 10^-3^ dilution rate. Plates were incubated at 30°C for 3 days and then imaged under blue light. Colonies showing GFP fluorescence in the uninduced samples are marked by white arrows. (D) PCR of a FACS-sorted GFP^+^ cell library to examine the gene rearrangements by SCRaMbLE in a synthetic *HIS* module. PCR primers are designed to target the *URA3* locus, within which the synthetic module is located.

### SCOUT enables rapid screening of SCRaMbLEd cells

Before applying SCRaMbLE to optimise synthetic *HIS* modules, we developed SCOUT (<u>S</u>CRaMbLE <u>C</u>ontinuous <u>O</u>utput and <u>U</u>niversal <u>T</u>racker), a fluorescent reporter system for tracking SCRaMbLE events based on a FLEX Cre-ON switch system^30^ (**Figure 2B**). In SCOUT, two pairs of heterotypic loxP-variant recombination sites, namely loxP2272 and loxP5171, are placed to flank a version of the *mGFPmut2* gene lacking the ATG start codon. The gene is placed in the antisense orientation with respect to a *TDH3* promoter with ATG start codon. Cre-mediated recombination between both pairs of the loxP recombination sites inverts the *mGFPmut2* gene in SCOUT and leading to a DNA construct giving constitutive GFP expression. This recombination also excises of one loxP-variant site from each pair, leading to stable and irreversible GFP expression. Cells exhibiting GFP fluorescence therefore are indicative of cells with nuclear Cre activity and thus are likely to be associated with SCRaMbLE events occurring in the genomes in those cells. SCOUT is designed as a plasmid-based system to allow us to transiently introduce it into strains ahead of inducing SCRaMbLE and then later remove it from screened strains by culturing in non-selective media. With this capability, SCOUT can be applied in iterative cycles of SCRaMbLE induction and screening.

The SCOUT plasmid was constructed using Golden Gate assembly and made compatible with the YTK^10^ and MYT^11^ systems, to allow for easy swapping of DNA parts such as promoters and selectable markers. We initially used a promoter (*pSCW11*) chosen to give SCRaMbLE events exclusively in daughter cells^31^. However, due to the reported leakiness of Cre-mediated recombination when using this strong promoter^19,22,25^, we then switched to the weaker *RET2* promoter as this gave reduced leaky expression and the cells still fully converted to GFP^+^ over time (**Figure S5**). To confirm whether SCOUT functions as expected, we induced SCRaMbLE in yeast with a defragmented *HIS* module and analysed the phenotypes and genotypes of the post-SCRaMbLE libraries after 4 hours of induction. We first confirmed reporter function by screening GFP fluorescent colonies, showing again the reduced leakiness of the *RET*2-based version (**Figure 2C**). Genotyping four randomly selected GFP^+^ colonies revealed SCRaMbLE-mediated gene rearrangements in all cases (**Figure S6**).

With SCOUT we can easily separate SCRaMbLEd cells (GFP^+^) from those without DNA rearrangements using fluorescence-based sorting (FACS). This sorting leaves us with a library of yeast cells with SCRaMbLE events highly likely in synthetic genome modules or other loxPsym-containing regions. When the regions of the genome containing these loxPsym sites are shorter than 35 kb we can use our recently described POLAR-seq method^26^ to determine the genotypes of the SCRaMbLEd regions using long-read sequencing and even apply this to the entire post-sorted library.

To determine if such an approach would work for our synthetic *HIS* modules, we took one of the module-containing yeast strains constructed for Figure 1, induced SCRaMbLE in a pool of these cells for 4 hours in the presence of the SCOUT construct and then used FACS to sort the population for GFP^+^ cells. We then extracted high molecular weight DNA from the post-FACS pool of SCRaMbLEd yeast cells and optimised long range PCR conditions to ensure we could get amplicons from this DNA pool that span the full synthetic *HIS* modules (**Figures S7A-C**). When SCRaMbLE and FACS were done in rich media conditions with histidine provided (+His) there was no selective pressure for the yeast to maintain the *HIS* genes. As expected, we saw diverse amplicon sizes arise from the PCR amplification from this yeast library, consistent with the variety of DNA rearrangements that SCRaMbLE provides, including deletion genotypes generated by SCRaMbLE in our library of cells (**Figure 2D**). In contrast, when the same experiment was done in media lacking histidine (-His) we observed uniform amplicons around 20 kb in length consistent with no deletions of the *HIS* genes in conditions where they are essential (**Figure 2D**). These positive PCR results show that post-SCRaMbLE libraries of synthetic *HIS* module containing yeast cells are suitable for POLAR-seq.

### SCRaMbLE and selection enriches for gene duplication in a sub-optimal module

Having established that SCOUT and FACS can generate high quality libraries suitable of POLAR-seq, we next examined SCRaMbLE rearrangements that can improve a poorly designed synthetic genome module. For this we focused on the *HIS*_refactor-4_ yeast strain that exhibited a major growth defect in the absence of histidine due to the mixed promoter strength refactoring (strain yXL219). The strain was transformed with the SCOUT plasmid (pXL005) and then SCRaMbLE was induced by adding 1 μM β-estradiol. After 4 hours of induction, over 1 million GFP^+^ cells were sorted by FACS (**Figure S8**) and a subset of these cells was then cultured in SC-His selective media to enrich for phenotypes that can grow well in the absence of histidine (**Figure 3A**). We isolated whole genomic DNA from this culture (called ‘yXL219_pool_’), amplified the DNA at the *URA3* locus by long range PCR and then sequenced the resulting pool of amplicons following the POLAR-seq method (**Figure S7A**).

**Figure 3.**
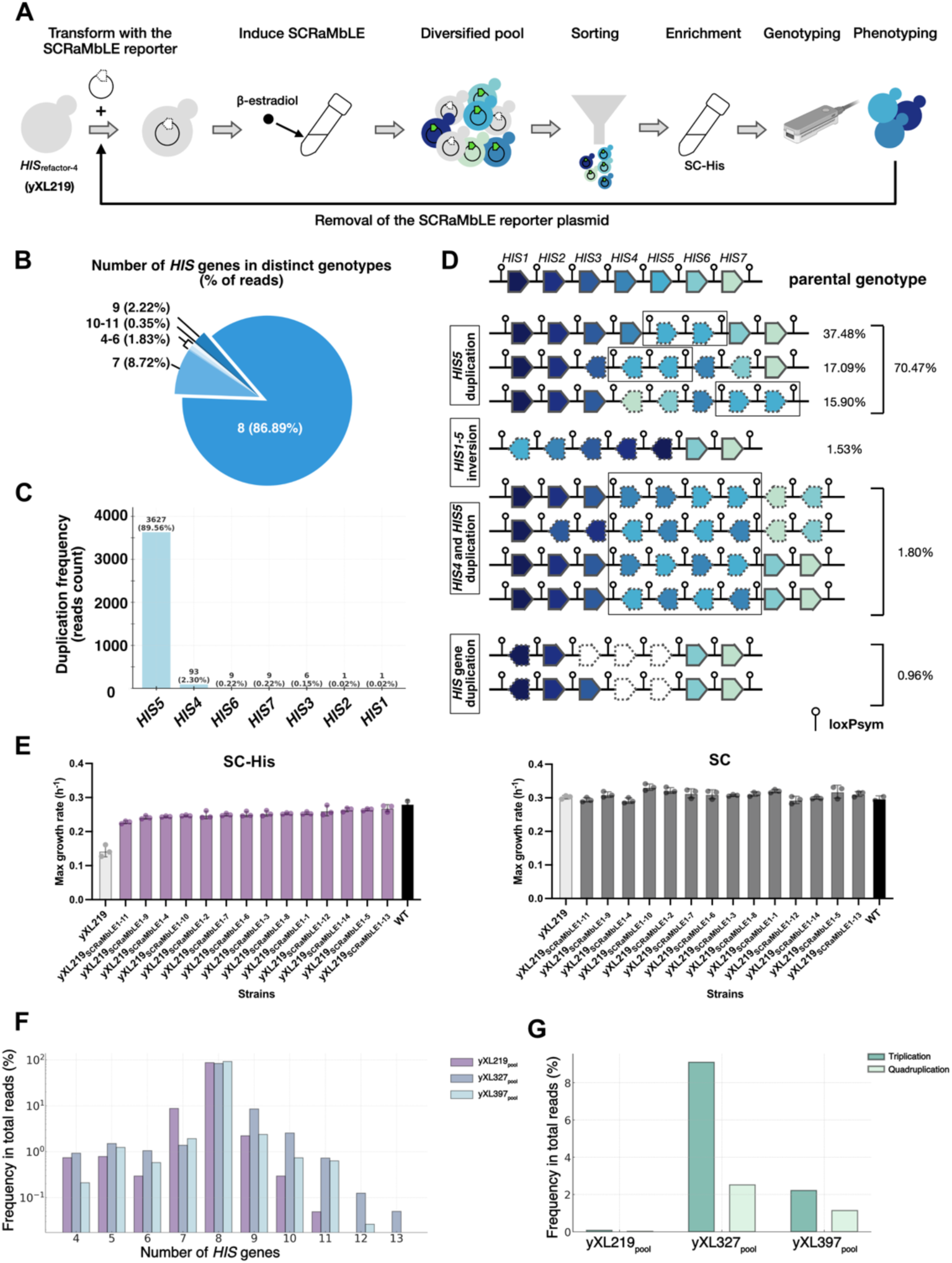
SCRaMbLE rescues function by duplication of *HIS5*. (A) Schematic of SCRaMbLE workflow for synthetic modules. Strain yXL219 that harbours the synthetic *HIS* module (*HIS*refactor-4) was transformed with a SCRaMbLE reporter plasmid (pXL005) which can induce Cre expression and indicate SCRaMbLEd cells by GFP fluorescence. Cells showing GFP expression were collected through FACS and subsequently grown in SC-His to enrich improved phenotypes. Genotyping of the post-SCRaMbLE library was performed through POLAR-seq. Top candidates with improved phenotypes were subjected to iterative rounds of SCRaMbLE after curing and re-introducing the SCRaMbLE reporter plasmid (pXL005). (B) Frequency of reads with different numbers of *HIS* genes, identified from the post-SCRaMbLE library yXL219pool. (C) Bar chart showing the duplication frequency of each gene in the detected reads from yXL219pool. (D) Schematic of representative post-SCRaMbLE genotypes identified in more than 10 reads from yXL219pool. Frequency in total reads is labelled on the right in percentage. Rearranged genes are highlighted with dashed lines. Gene duplications are highlighted with black squares. (E) Maximum growth rates of the 14 strains isolated from yXL219pool. Bars in grey, purple, and black represents the defective parental strain yXL219, the strains isolated from yXL219pool and the WT control strain (yXL014), respectively. Cultures were grown at 30°C and performed in 3 biological replicates. Error bars represent standard deviation. Culture medium is stated above each graph. (F) Bar chart showing the numbers of *HIS* genes in each distinct genotype identified from yXL219pool, yXL327pool and yXL397pool and their frequency in total reads. (G) Frequency of reads detected with *HIS* gene triplication or quadruplication from yXL219pool, yXL327pool and yXL397pool.

POLAR-seq identified 127 distinct genotypes from 4,050 annotated full length reads (**Figure S7D**). Approximately 86.89% of the total reads displayed genotypes containing 8 transcription units (TUs) within the synthetic *HIS* module, indicating duplication of one gene (**Figure 3B**). A small portion of reads (2.57%) had 9 or more TUs. Duplication of *HIS5* was the overwhelmingly dominant trend with it being evident in nearly 90% of all reads (**Figure 3C**). The fact that rearrangement and then growth in histidine lacking media selects for cells with duplicated *HIS5* is consistent with our earlier hypothesis that poor *HIS*_refactor-4_ growth was due to insufficient expression of *HIS5* in this strain.

As well as revealing the abundant genotypes with *HIS5* duplications, POLAR-seq also detected some rare genotypes that would likely be challenging to identify by colony screening methods. Four distinct genotypes, each including two copies of both *HIS4* and *HIS5* TUs, comprised 1.80% of the total reads (**Figure 3D**). We also identified two deletion genotypes, constituting 0.96% of the total reads. The detection of deletion-containing module genotypes in cells grown in media lacking histidine is surprising. This may just be amplification of modules from dead or non-growing post-SCRaMbLE cells. In general, it is important to note that the percentage of each genotype determined by POLAR-seq might not precisely reflect the actual sub-population composition in a population of cells, as there may be enrichment and more sequencing of shorter amplicons, for example due to PCR bias^32^.

To confirm whether the main enriched genotypes from POLAR-seq correlate with improved histidine biosynthesis, we picked 14 GFP^+^ colonies at random from the same FACS-sorted library and characterised their growth fitness in SC-His and SC media conditions. We observed a significant increase in the growth rates of all 14 strains in SC-His compared to the parental strain yXL219 (**Figure 3E**) demonstrating the fitness improvement post-SCRaMbLE. Strain yXL219_SCRaMbLE1-13_ exhibited the highest maximum growth rate, almost close to that of the WT control. When histidine was added to the growth media, growth rates were similar across all tested strains (**Figure 3E**).

Clonal long-read sequencing of the module region for 8 out of these 14 strains revealed a spread of genotypes that aligns well with the POLAR-seq data described above. Notably, the genotypes of 7 of the 8 sequenced strains were among the top three most abundant genotypes identified by POLAR-seq, and all exhibited *HIS5* duplication (**Figure S9A**). From these results, we confirm that genotype enrichment is indeed likely to be associated with phenotypic improvement in histidine biosynthesis and that the POLAR-seq method accurately captures this.

### A second cycle of SCRaMbLE has limited scope for module improvement

Although a single round of SCRaMbLE may generate significantly improved phenotypes, we have previously seen in other work that function-related phenotypes such as pigment production, can be cumulatively improved through iterative SCRaMbLE^21,22,33^. Thus, to explore the possibility of further improving growth fitness of the strains isolated from yXL219_pool_, we performed a second round of SCRaMbLE on two strains that showed promising results in the first round. These two strains, namely yXL219_SCRaMbLE1-2_ and yXL219_SCRaMbLE1-13,_ were chosen because they were the most enriched genotype (**Figure S9A**) and the fastest growth phenotype (**Figure 3E**), respectively. Before starting the second round, the SCRaMbLE reporter plasmid was removed from both strains, and reintroduced fresh through re-transformation (**Figure 3A**). The second round of SCRaMbLE was carried out under the same conditions as the first round, generating two post-SCRaMbLE libraries, designated as yXL327_pool_ for yXL219_SCRaMbLE1-2_, and yXL397_pool_ for yXL219_SCRaMbLE1-13_.

In POLAR-seq analysis of the post-SCRaMbLE libraries yXL327_pool_ and yXL397_pool_, we found that the most abundant genotypes in these two libraries kept the genotype of their parental strains yXL219_SCRaMbLE1-2_ and yXL219_SCRaMbLE1-13_, respectively (**Figure S9B and S9C**). This suggests that the growth advantage improvements in strain yXL219_SCRaMbLE1-2_ and yXL219_SCRaMbLE1-13_ may have already approached a local optimum in histidine lacking media. In addition, the significantly lower proportion of other genotypes in these cell libraries makes it less efficient to screen colonies showing the non-parental genotype (**Figure S9B and S9C**). Based on the lack of growth improvement we halted the iterative SCRaMbLE after just the two cycles.

We next examined the size variations of the synthetic *HIS* module in our second-round SCRaMbLE libraries. Eight-gene modules with *HIS5* duplication still dominate the population after this round (**Figure 3F and S10**). However, we noted a few cases of further expansion in the synthetic *HIS* module, with increased presence of amplicon reads with 11, 12 and 13 genes in the module (**Figure 3F**). The largest module we identified had duplications of *HIS2*, *HIS3*, *HIS4* and a quadruplicate of the *HIS5*, with a theoretical module size of ∼35 kb. We also observed a significant increase in genotypes showing triplication or quadruplication of *HIS5* in the second-round libraries (**Figure 3G**). This suggests a continuing enrichment of cells with more copies of the *HIS5* gene when grown under histidine selective conditions. Further iterative SCRaMbLE of these genotypes would be interesting to investigate, but unlikely to give significant growth improvements. Overall, both SCRaMbLE and iterative SCRaMbLE on the yXL219 strain revealed that the *HIS*_refactor-4_ module would benefit from being redesigned to have double the expression of *HIS5* (i.e. by use of a stronger promoter for this gene) but that further design changes, such as increased expression of other genes or alternative gene locations and directions would not make any significant difference to function.

### Iterative SCRaMbLE in a synthetic chromosome to maximise phenotype

Having observed SCRaMbLE achieving a solution to poor genome module design in just a single cycle, we wanted to explore whether this was due to the nature of the phenotype we were selecting for or due to the limited possibilities for viable gene rearrangements in the *HIS* module. Presumably if SCRaMbLE is genome- or chromosome-wide there are many more viable rearrangements possible and so it may take more cycles to reach phenotypic improvements. To test this, we performed SCRaMbLE across SynV, a whole synthetic chromosome produced for the Sc2.0 project that replaces wildtype chromosome 5 of the *S. cerevisiae* genome. We chose to screen and select for a simple phenotype that doesn’t have an obvious solution, as it did in the *HIS* module case. We screened and selected for maximising green fluorescence per cell, when the strain is constitutively expressing sfGFP. To avoid a simple genetic solution, like duplication of the sfGFP gene, we encoded the sfGFP gene on a yeast plasmid (pBAB016) without any loxPsym sites, so it cannot SCRaMbLE. The strain created for this study, synV-pBAB016, is a haploid.

To begin SCRaMbLE, synV-pBAB016 was transformed with the Cre recombinase expressing plasmid *pSCW11*-creEBD. SCRaMbLE was then induced with 1 μM β-estradiol for 4 hours, after which time cells were washed twice and plated at various dilutions. A control culture was included that was not induced. After 2 days growth on plates, 3 uninduced control colonies were picked randomly alongside 36 of the brightest post-SCRaMbLE colonies. These were selected by eye under far blue light illumination, with all colonies picked into SC-His-Leu media. Cultures were grown overnight and back diluted 1:100 into fresh SC-His-Leu media in a 96-well plate. Endpoint GFP fluorescence per well was then measured after 48 hours of growth. The strain with the highest OD_600_-normalised fluorescence (a proxy for GFP expression per cell) was isolated. This top performing strain in the first round of SCRaMbLE was termed ‘R1’. This strain was next subject to a second round of SCRaMbLE. This process was repeated five times, resulting in strains R0 (pre-SCRaMbLE), R1, R2, R3, R4, and R5 (**Figure 4A**).

**Figure 4.**
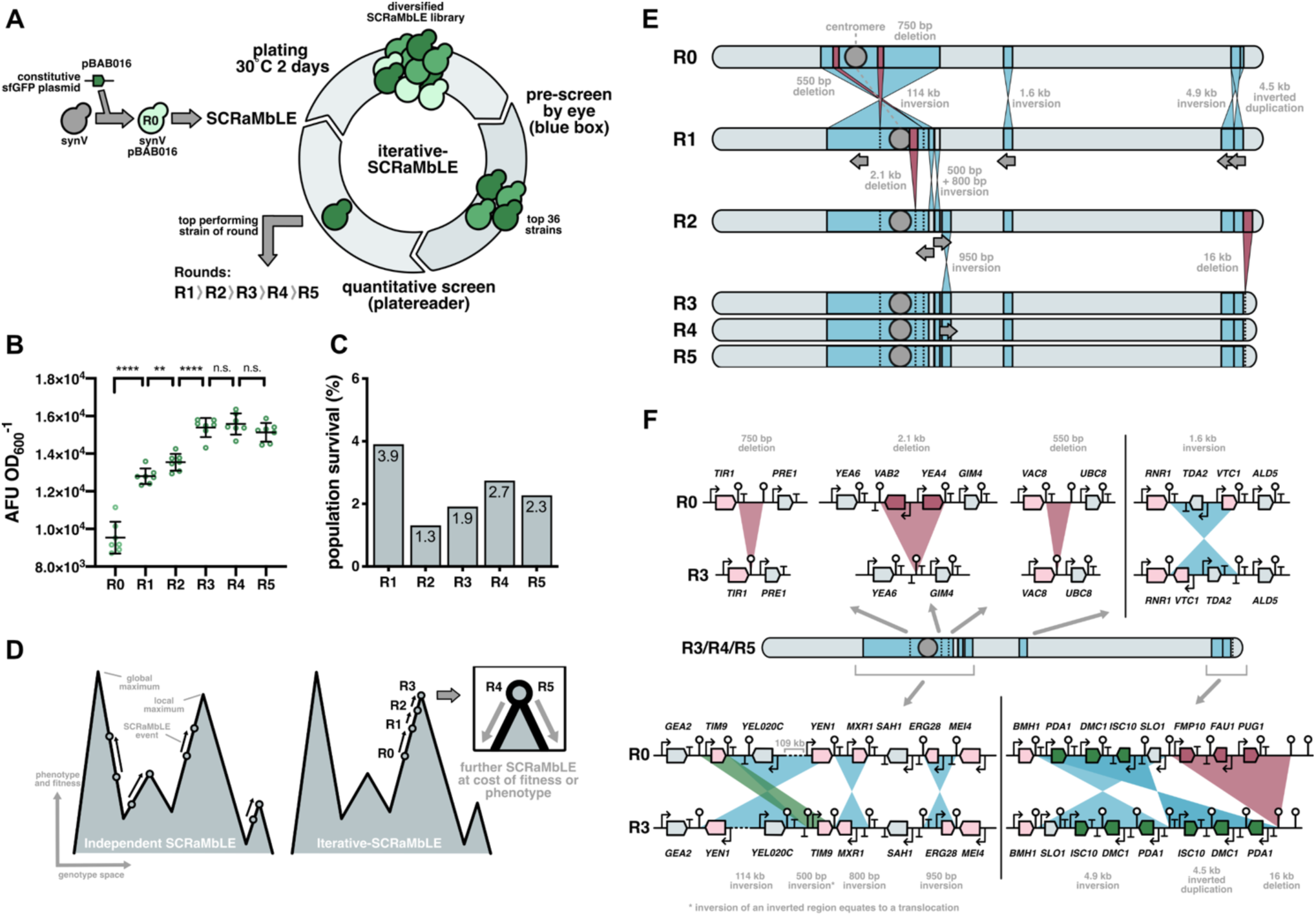
Increasing sfGFP expression by iterative SCRaMbLE of a synthetic chromosome. (A) SCRaMbLE was induced in strain synV-pBAB016-*pSCW11*-creEBD (R0) for four hours to create a diversified SCRaMbLE library. The top 36 strains were selected, and fluorescence characterised using a plate reader. The top performing strain (R1) was stocked and subsequently subject to a second round of SCRaMbLE, as before, to produce strain R2, this was repeated for strains R3, R4, and R5. (B) Strains were selected and picked by eye under illumination of blue light (‘blue box’). Endpoint 520 nm fluorescence and OD600 was measured after 48-hour growth. Mean is shown with error bars that indicate standard deviation for n=7 samples. ****, p<0.0001; **, p<0.01; n.s., not significant. (C) Colonies were counted for induced and uninduced cultures after each round of SCRaMbLE (R1-R5). Population survival is shown as the percentage of induced colonies to uninduced colonies. For clarity values for each bar are shown as numbers. (D) A simplified two-dimensional depiction of a phenotype and fitness landscape. Transition of a cell (circle) upwards indicates an improvement in fitness and/or phenotype. Arrows indicate individual rounds of SCRaMbLE. During independent SCRaMbLE experiments many local maxima are sampled but not necessarily reaching the optimum fitness/phenotype within each maximum. Iterative-SCRaMbLE commits a cell lineage to a single local maximum. Once at the optimum fitness/phenotype (depicted here as occurring in R3) no further fitness or phenotype improvements can be seen with subsequent rounds of SCRaMbLE (R4 and R5, inset). Any further SCRaMbLE events, in combination with those already existing result in decreased fitness and/or phenotype. (E) Rearrangement events at each round of iterative SCRaMbLE determined by multiplexed long-read sequencing. Inversions are shown in blue. Grey arrows below each chromosome indicates the new orientation of the region. Deletions are shown in red with the affected region subsequently marked with a dotted vertical line. The centromere is depicted as a grey circle. (F) Summary of SCRaMbLE events in strain R3 (and R4-R5). Deletions are shown in red, inversions are shown in blue, and translocations are shown in green. Pink genes indicate those with a disrupted 3’ UTR, red genes indicate those that are deleted, and green genes indicate those that have undergone a copy number change through a duplication event. LoxPsym sites are indicated by white circles on sticks.

To directly compare the sfGFP expression of each of these strains, we first removed *pSCW11*-creEBD plasmid from the strains R0-R5 by passaging through two overnight cultures in SC-His media. After confirming the loss of Cre plasmid, we randomly picked 7 colonies for each strain and measured the 48-hour endpoint OD_600_ and GFP fluorescence. We observed a significant increase in sfGFP fluorescence within the first 3 rounds of SCRaMbLE (R0-R3). However, the phenotype did not significantly improve in the fourth and fifth round (R4 and R5) (**Figure 4B**). This indicates that either no new SCRaMbLE events could yield further phenotype improvement or that no additional SCRaMbLE events occurred in the final 2 rounds.

We monitored cell viability by counting colonies for both the uninduced and induced cultures in each round of SCRaMbLE. We calculated the population survival by the percentage of induced colonies compared to uninduced. Population survival was highest for R1 (3.9%) and was lower for all subsequent rounds of SCRaMbLE, suggesting that the presence of existing SCRaMbLE events increased the sensitivity of cell viability to additional SCRaMbLE events (**Figure 4C**). This presumably puts an upper limit on the number of SCRaMbLE events that can be accumulated before the cell is non-viable, representing a local maximum in the phenotype-fitness design (**Figure 4D**).

To understand how cumulative SCRaMbLE recombination events accumulate phenotype improvements, we sequenced the strains R1-R5 using multiplexed barcoded long-read sequencing. A total of eight samples including R1-R5 were sequenced on R9.4.1 flowcell in a MinION Mk1B device. Run statistics for samples R1-5 are shown in **Table S10**. By aligning contigs and corrected reads to a pre-SCRaMbLE scaffold sequence, we determined all SCRaMbLE events for each round of SCRaMbLE, as shown in **Figure 4E**. As expected, the first round of SCRaMbLE produced the greatest number of SCRaMbLE events, including the inversion of a 114 kb region, which led to the reposition of the centromere and contained two small deletions. R2 and R3 revealed less recombination events than R1. Interestingly, no additional SCRaMbLE events were detected in R4 and R5, suggesting any additional SCRaMbLE events in combination with those present in R3, could not improve the phenotype any further without the loss of cell viability.

By comparing the final chromosome rearrangement (R3-R5) to the pre-SCRaMbLEd chromosome (R0), we were able to characterise exactly which genes were affected by each recombination event, as shown in **Figure 4F**. Aside from potentially PDA1, which was inverted and duplicated when generating R1 and encodes an enzyme that affects carbon supply to the TCA cycle, none of the genes that were affected by any of the SCRaMbLE events observed have an obvious link to sfGFP protein production. Interestingly, none of these recombination events were seen in another previously described strain called VB2 that similarly underwent SCRaMbLE and showed enhanced GFP fluorescence^34^. This is again consistent with our assumption that the phenotype-fitness landscape is rugged and different improved phenotypes (local maxima) can be achieved through independent SCRaMbLE routes (**Figure 4D**).

## Discussion

In this work, we developed a new tool, SCOUT, and described methods for iterative SCRaMbLE to be used to improve desired phenotypes in yeast strains with synthetic genome modules and synthetic chromosomes. To provide a testbed for this work, we also described reorganisation of the 7 genes of the yeast histidine biosynthesis pathway into different designs of synthetic genome modules, including both ‘defragmented’ and ‘refactored’ versions. Notably, no obvious fitness defects were observed when defragmenting these genes or when refactoring the genes by using three sets of synthetic constitutive promoters with similar strengths, presumably with all these promoters lacking the native regulation of the *HIS* gene promoters. These results confirmed either the robustness of the histidine biosynthesis pathway to alternative levels of gene mRNAs or robustness of the yeast host itself, in terms of being able to tolerate a wide range of *HIS* enzyme expression levels. Naturally, the yeast histidine biosynthesis pathway is regulated at the protein level by end-product feedback inhibition, and this may explain why removing native promoters, and thus transcriptional regulation, does not have a major effect on pathway function. Transcriptional regulation of these genes is known to involve the general transcription factor GCN4, transcription factors Bas1 and Bas2(Pho2) and histidine biosynthesis is also co-regulated through transcriptional regulation and via intermediate AICAR with purine biosynthesis pathway^35–39^. However, in the limited conditions we assessed cell performance here, we saw no evidence that this native transcriptional regulation was required. Instead, we speculate that in YPD, SC and SC-His media metabolic flux toward histidine biosynthesis and related pathways is likely kept at a functional basal level regardless of *HIS* promoter-encoded features.

While this work and that of others has shown the tolerance of yeast genes to changes in sequence, location and even entire promoters, it is important to note that relocating and refactoring genes may impair cell viability, especially if the genes in question encode an essential function^15,16^. Due to this risk, we propose that synthetic genome module construction should be designed to incorporate some form of *in vivo* diversification system, such as the SCRaMbLE system used here. This then allows poor performing constructs to quickly be debugged via diversification and growth- or function-linked selection. Similar to our approach, a recent study employed a variant of SCRaMbLE, named GEMbLeR, but in this case, specifically designed to shuffle only promoters and terminators using orthogonal LoxPsym sites, aiming to diversify gene expression and optimize heterologous pathway function^40,41^. Compared with mutagenesis-based approaches, such as adaptive laboratory evolution (ALE)^42^, OrthoRep^43,44^ and yEvolvR^45^, the SCRaMbLE approach offers an alternative that focuses on adjusting gene expression by changing the copy numbers or orientations of modular DNA regions in the genome. The downstream analysis of the genomic changes leading to functional improvements is therefore more straightforward, as SCRaMbLE only involves region-specific rearrangements, whereas other methods generate more classical mutations like SNPs and Indels that can be harder to identify (especially when genome-wide) and are challenging to interpret phenotypic improvements from. Determining the genotype changes from SCRaMbLE has become relatively straightforward in the last decade with the emergence of long-read DNA sequencing, including targeted methods, like POLAR-seq, that can compare hundreds of SCRaMbLE outcomes at low cost^26^. If duplications of genes within a SCRaMbLEd module are seen as clearly beneficial (as seen in the *HIS*_refactor-4_ case here), then it informs that a stronger promoter is needed for the genes that are duplicated. Conversely, if genes are lost, then it identifies that they are not needed for the module’s function in the assayed conditions. The use of iterative cycles of SCRaMbLE also provides a way to determine whether further module design changes can give significant improvements to module function and cell performance.

Prior work has shown that iterative SCRaMbLE can accumulate phenotype improvements in yeast, such as in heterologous β-carotene or prodeoxyviolacein production^21,22,33^. Here, we took this further and used long-read sequencing over several SCRaMbLE cycles to investigate how cumulative genotype changes led to phenotype improvements from strains with either synthetic modules or entire synthetic chromosomes. In both cases, we observed that the first round of SCRaMbLE yielded strains with notable phenotypical improvements, but subsequent rounds displayed a rapid plateau, indicated by the stable recurrence of the predominant genotype. This is consistent with iterative-SCRaMbLE committing to a single local maximum in the phenotype-fitness landscape. It’s important to note, however, that this maximum may not be the global optimum and that it could be limited by the scope of the rearrangements done in the first round. With this in mind, we recommend that the first round of SCRaMbLE is scaled to achieve maximum diversity, for example by rearranging and screening/selecting with a large population. Our development of the SCOUT reporter and the use of this with the POLAR-seq method offers the ideal tools for using SCRaMbLE on larger populations. SCOUT combined with FACS enables users to rapidly isolate the population subset likely to have the most SCRaMbLE events as a pool and so aids in enriching the screening/selection step for the most diverse outcomes. This works especially well when SCRaMbLE is being used on a synthetic region short enough to generate full length amplicons for POLAR-seq, so that the genotypes in the pool can be sequenced and compared in bulk.

Together the work described here demonstrates the combinatorial possibilities afforded by synthetic biology when redesigning genomic regions and offers new tools and methods to effectively navigate this. Using iterative SCRaMbLE, we have shown that rapid, targeted modular genetic modifications can rescue the functionality of defective synthetic modules and quickly find genotype solutions to phenotype improvements on a chromosome-wide scale. Our study also demonstrates how genome refactoring can be done at the module scale and still achieve efficient function for a key metabolic pathway, and it shows how the approach can efficiently obtain genotype-to-phenotype datasets from POLAR-seq that are valuable for determining ideal gene expression profiles for cells grown in different conditions. We anticipate that datasets like these will be crucial for the future of designing custom modular synthetic genomes.

## Methods

### Strains and media

Yeast strains generated in this study are derived from BY4741 yeast (*<u>MATa</u> his3Δ1 leu2Δ0 met15Δ0 ura3Δ0*)^27^, including the Sc2.0 project strain synV-pBAB016^34^. See **Supplementary Table 1** for a full list of derived yeast strains. NEB Turbo competent *Escherichia coli* (*E. coli*) from New England Biolabs (NEB) was used for all DNA cloning and plasmid propagation work.

<u>Y</u>east extract <u>P</u>eptone Dextrose (YPD) media (10 g L^-1^ yeast extract (VWR), 20 g L^-1^ peptone (VWR), 20 g L^-1^ glucose (VWR)) was used for general culturing of yeast cells, unless otherwise stated. <u>S</u>ynthetic <u>C</u>omplete media (SC; 6.7 g L^-1^ Yeast Nitrogen Base without amino acids, 1.4 g L^-1^ Yeast Synthetic Drop-out Medium Supplements without L-uracil, L-tryptophan, L-histidine, L-leucine, 20 g L^-1^ glucose) was used for auxotrophic selection experiments, or was used with all amino acids supplemented as a defined complete medium. Amino acids such as 20 mg L^-1^ L-tryptophan, 20 mg L^-1^ L-histidine, 20 mg L^-1^ uracil and 120 mg L^-1^ L-leucine were supplemented into SC media depending on the required auxotrophic selection. For growth on plates, media were supplemented with 20 g L^-1^ bacto-agar (VWR). Unless otherwise stated, all other components used in the media were supplied by Sigma Aldrich.

Luria-Bertani (LB) medium was used for culturing *E. coli*. LB agar was prepared by dissolving 37 g L^-1^ LB agar powder (VWR) into required amount of distilled water. Antibiotics such as ampicillin (100 μg mL^-1^), chloramphenicol (34 μg mL^-1^), kanamycin (50 μg mL^-1^) and spectinomycin (100 μg mL^-1^) were supplemented when necessary.

### Plasmids

A full list of plasmids generated in this study can be found in **Supplementary Table 2** (gap repair donor and gene fragment plasmids)**, Supplementary Table 3** (linker plasmids), and **Supplementary Table 7** (pre-assembled linker vectors). Unless otherwise stated, all plasmids were constructed using the MoClo Yeast Toolkit (YTK)^10^. Standard parts from the YTK libraries, such as promoters, terminators, and plasmid backbones, were typically used, with these stored in entry vectors by the group. For parts not found in the group’s standard libraries, DNA was typically amplified by PCR using primers that add the necessary YTK-compatible overhangs so that the PCR product can be directly used in its intended YTK assembly. A few parts were kindly provided by other Ellis Lab members, see acknowledgements in the parts list (**Supplementary Table 6**). These parts were then assembled into cassettes (listed in **Supplementary Table 8**) and the cassettes were then assembled into multigene cassettes (listed in **Supplementary Table 9**) through the standard YTK assembly workflow^10^. For a full list of oligos and their sequence, see in **Supplementary Table 4**.

Golden Gate Assembly was used to assemble plasmids whenever the DNA sequences being assembled are free of recognition sites for the type IIs restriction enzymes used by the Golden Gate reaction, i.e. BsmBI, BbsI, or BsaI. Gibson assembly was used when the sequence of DNA fragment to be assembled was not free of recognition sites of the type IIs restriction enzymes BsmBI, BbsI, and BsaI.

### Genomic DNA isolation for PCR

Genomic DNA of yeast was isolated following a LiOAc/SDS isolation protocol^46^. Plasmids were isolated the following Qiagen Miniprep protocol. Q5 high-fidelity DNA polymerase (NEB) PCR conditions and protocols were used for the DNA amplification from the extracted genomic DNA.

### Yeast transformation

A colony was picked out from the plate and grown to saturation in 2 mL appropriate media overnight (30°C, 250 rpm). The next day, cell culture was diluted to OD_600_ ∼0.2 in a 50 mL Falcon centrifuge tube with 10 mL fresh media and grown for ∼6 h to OD_600_ = 0.8-1.0. Cells were pelleted by centrifuging at room temperature (2000 rcf, 10 minutes), then washed once with 10 mL 100 mM lithium acetate (LiOAc, Sigma Aldrich). Centrifugation was repeated and cell pellet was resuspended in ∼600 μL 100 mM LiOAc. 100 μL of this mixture was added to 64 μL DNA cocktail containing 10 μL of boiled salmon sperm DNA (ThermoFisher) per transformation, and then gently mixed with 296 μL PEG-3350/LiOAc mixture (260 μL 50% (w/v) PEG-3350 and 36 μL 1M LiOAc). This mixture was then placed in a heat block at 42°C for 40 min and cells were pelleted by centrifugation at 8000 rpm for 1 min. Pellets were resuspended in 100-200 μL 5 mM CaCl_2_. After a 10 min recovery, cells were plated onto the appropriate agar media for selection.

### CRISPR/Cas9 genome engineering

For multiplex gene deletion, 50 ng of the CRISPR/Cas plasmid (pWS2081-*URA3*), 600 ng of each sgRNA plasmid were mixed together with 0.5 μL BpiI (ThermoFisher), 1 μL of 10X Buffer G (ThermoFisher), and nuclease-free water to make up to 10 μL. This mixture was then incubated at 37°C for 8 hours followed by 80°C heat inactivation for 10 min. 5 μg of each donor DNA was added to this mixture to a total volume of 64 μL to be used for the yeast transformation. Donor DNA was generated by PCR amplification from the genomic DNA and purified by DNA Clean & Concentrator Kit (Zymo Research). gRNA plasmids were constructed by T4 PNK phosphorylating (NEB) and annealing two oligos, followed by a BsmBI Golden Gate assembly to insert the small fragment into the SpCas9 sgRNA Dropout vector (Ellis lab plasmid pWS2069). Oligos for gRNAs were designed by adding the sequence 5’-AGAT-3’ at the 5’ end of the guide sequence and 5’-AAAC-3’ at the 5’ end of the reverse complementary of the guide sequence.

Unless otherwise stated, for genome integration and other genome editing experiments, 250 ng of the CRISPR/Cas plasmid and 500 ng of each DNA fragment was combined with 10 μL boiled salmon sperm DNA, made up to 64 μL with nuclease-free water to be used for the yeast transformation. DNA fragment was generated by PCR amplification and purified by DNA Clean & Concentrator Kit (Zymo Research). gRNA plasmids were constructed according to the ‘gRNA-tRNA Array Assembly’ methods described in the Multiplex MoClo Toolkit^11^.

For a full list of gRNAs used for CRISPR/Cas9 genome engineering, see **Supplementary Table 5**.

### Plasmid curing

For curing *URA3* containing plasmids, 5-FOA (5-fluoroorotic acid) counterselection was used^47^. Colonies were inoculated into 2 mL YPD media and grown overnight (30°C, 250 rpm). This culture was streaked using a 10 μL loop onto the agar plate supplemented with 5-FOA (Formedium). Colonies showing growth after incubation for 3 days at 30°C suggested successful plasmid curing.

For an auxotrophic marker that is not *URA3*, strains were firstly streaked onto the YPD agar plate. After 3 days of incubation at 30°C, colonies were picked and inoculated into 2 mL of fresh YPD media and grown overnight (30°C, 250 rpm). This culture was then streaked using a 10 μL loop onto the YPD agar plate and incubated at 30°C for 3 days. Colonies were then streaked onto both the agar media plates with selection and also agar media plates lacking selection. Strains showing growth on the media lacking selection but not the media with selection suggested successful plasmid curing.

### SCRaMbLE

Strains transformed with either pXL004, pXL005 (SCRaMbLE reporter plasmids) or plasmid *pSCW11*-creEBD were grown overnight in 2 mL of appropriate media (30°C, 250 rpm). This culture was diluted in 5 mL of fresh media to OD_600_ ∼0.2 and grown for ∼4 h to OD_600_ = 0.4-0.5. SCRaMbLE was induced by adding 1 μL of 5 mM β-estradiol dissolved in DMSO to the culture. For the control (uninduced), 1 μL of DMSO was added. Cultures were grown for an additional 4 h before being washed twice in water and resuspended in 1 mL PBS. Cells were diluted x10^-3^ and x10^-4^ and plated onto appropriate agar plates and grown for 2-3 days at 30°C.

### Iterative-SCRaMbLE

For the iterative SCRaMbLE of synthetic *HIS* modules, after each round of SCRaMbLE and POLAR-seq, the phenotypes and genotypes of several of the most abundant strains were characterised individually. The top performers were then grown in rich media to remove the SCRaMbLE reporter plasmid (pXL005), in which the sfGFP gene is reversed. The strains confirmed with the removal of this plasmid were then reintroduced with a new pXL005 and subjected to the next round of SCRaMbLE.

After each SCRaMbLE round of the strain synV-pBAB016, 36 SCRaMbLE colonies and 3 control colonies were picked into 500 μL selective media in a 2.2 ml deep well 96-well plate (Grenier). After a single overnight growth (30°C, 250 rpm) 1μL was used to inoculate 99 μL of appropriate selective media and OD_600_ and endpoint fluorescence measured in a plate reader at 520 nm after 48 h of shaking growth at 30°C. The strain with the highest OD_600_-normalised fluorescence was streaked out and used to inoculate 5 mL of fresh media for a subsequent round of SCRaMbLE as well as being stocked in 25% glycerol at -80°C. Once a high performing strain was obtained for each round of SCRaMbLE these top strains were streaked onto appropriate selective media lacking selection for curing the Cre-recombinase plasmid.

### Fluorescence-Activated Cell Sorting (FACS)

FACS sorting was performed on the BD FACSAria III Cell Sorter (BD Biosciences) to select yeast cells with GFP fluorescence. Yeast cells, washed and resuspended in PBS buffer post-SCRaMbLE, were transferred to a 5 mL FACS tube (Invitrogen) and diluted to appropriate density with PBS buffer for FACS sorting. The 70 μm nozzle was selected for the sorting. GFP^+^ cells sorted from the FACS instrument were collected in a 15 mL Falcon centrifuge tube. A subset of cells was immediately inoculated into SC-His and grown for 2-3 days to reach saturation (30°C, 250 rpm). The remaining cells were spun down at 4000 rpm for 20 min. The pellet was resuspended in 0.25 mL PBS and stocked in 25% glycerol at -80°C.

For the FACS analysis, yeast cells were firstly selected based on morphology (FSC-A vs SSC-A). Single cells were then selected based on a double doublet-discrimination (FSC-A vs FSC-H and SSC-W vs SSC-H). Single GFP^+^ cells were then selected based on GFP expression (FSC-A vs GFP). GFP expression was selected with the 530/30 bandpass filter.

### High Molecular Weight (HMW) genomic DNA isolation

HMW genomic DNA of the post-SCRaMbLE cell libraries was isolated according to the method published by Denis *et al*.^48^ with the following modifications in the protocol: yeast culture was grown in 50 mL of appropriate media overnight until OD_600_ = 5-15, Zymolyase (Zymo Research) was replaced with Lyticase (Sigma Aldrich, 600 U per 1 mL of OD_600_ = 1), and all centrifugation steps were performed at 4000 g. HMW genomic DNA of the individual strains was isolated from 2 mL of overnight yeast culture using the adapted protocol and proportionally reduced reagents.

HMW genomic DNA of the post-SCRaMbLE synV-pBAB016 strains R1-R5 was isolated for nanopore sequencing using Genomic-tip Kits (Qiagen) according to the instructions from the manufacturers. Any handling of DNA was done using wide bore tips where appropriate.

### POLAR-seq

POLAR-seq and data analysis were performed according to the method published by Ciurkot *et al*^26^.

Long range PCR of the full synthetic *HIS* module was set up in a 25 μL of LA Taq Hot Start (TaKaRa, Shiga Japan) PCR reaction according to the manufacturers’ protocol, using 20 ng of genomic DNA as the template and primers at a final concentration of 0.1 μM. Amplicon was verified by gel electrophoresis (50 V for 5 h) on a 0.6% agarose gel.

The pool of amplicons was prepped for sequencing with NEBNext Companion Module and the ligation kit SQK-LSK109 or SQK-LSK114. The library was sequenced on Flongle (FLG001 or FLG114) using MinION Mk1B. The data was collected with the latest version of MinKnow (22.05.5 – 23.04.6).

### Nanopore sequencing of the post-SCRaMbLE synV-pBAB016 strains (R1-5)

Genomic DNA was quantified using a Qubit 2.0 fluorimeter to ensure a minimum amount of 1 μg present in at least 50 μL volume (20 ng/μL). The DNA library was prepared using the SQK-LSK109 kit with 1D native barcoding EXP-NBD104 kit (Oxford Nanopore Technologies). After ligation of the barcoding sequences BC01-05 to samples R1-R5, respectively, samples were pooled, along with 3 other barcoded samples. A total of eight samples including R1-R5 were sequenced on a single R9.4.1 flowcell in a MinION Mk1B device. Sequencing was allowed to proceed for 48 hours using the latest version of MinKnow software with local basecalling.

Raw reads were corrected using the correction step of canu v1.8 (www.canu.readthedocs. io) to create a set of long reads (N50 ∼45 kb) with a higher read accuracy and sequencing genome coverage of 40x. De novo contigs were generated using these corrected reads with smartdenovo (https://github.com/ruanjue/smartdenovo) and both contigs and corrected reads were aligned to the pre-SCRaMbLE synV sequence using LASTAL. Alignments were viewed in Tablet with a synV GFF3 file to identify recombination events with boundaries at loxP sites^49^. To better facilitate multiplexing in sequencing analysis each bioinformatic program was run through a bash shell script to automatically process R1-R5 reads sequentially.

### Plate reader assay

Overnight cultures were harvested, washed and used to inoculate 100 μL cultures in a 96-well plate with a starting OD_600_ normalised to 0.02. Plates were incubated and measured in a Synergy HT Microplate Reader (Biotek) shaking at 30°C. Mean absorbance values of equivalent blank media wells were subtracted from data points.

### Microscopy

A single yeast colony was used to inoculate 2 mL of appropriate media. Cultures were grown overnight (30°C, 250 rpm) and visualised on a Nikon Eclipse Ti inverted microscope at 20x magnification and optical conf. Bright field (BF) images were captured using the Nikon NIS-Elements Microscope Imaging Software. Fiji^50^ was used to process the images and add the scale bars.

## Acknowledgements

This research was supported by a Chinese Scholarship Council (CSC) PhD scholarship to X.L. and Wellcome Trust Discretionary Award (221267/Z/20/Z) providing funding for K.C. and T.E. W.M.S. and G-O.F.G. were supported by BBSRC CASE PhD studentships.

## Supplementary Figures

**Figure S1.**
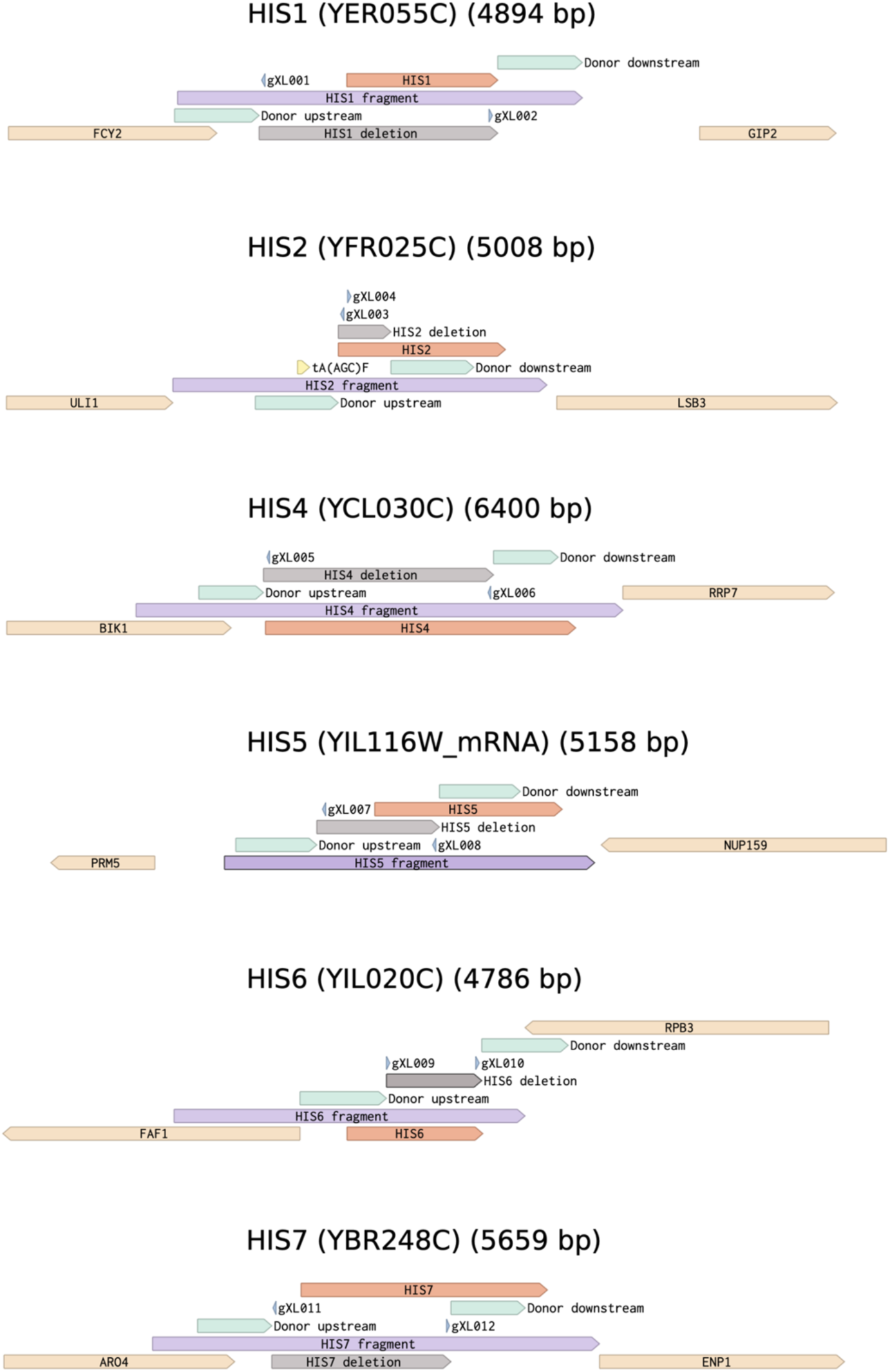
Schematic for gene deletion and defragmentation of *HIS* genes on Benchling (www.benchling.com). The CDS of genes are indicated by dark orange. Neighbouring genes are marked in light orange. Deletion regions are highlighted in grey. Gene fragments for defragmentation of the synthetic *HIS* module are shown in purple. tRNA gene tA(AGC)F upstream of *HIS2* is marked in yellow.

**Figure S2.**
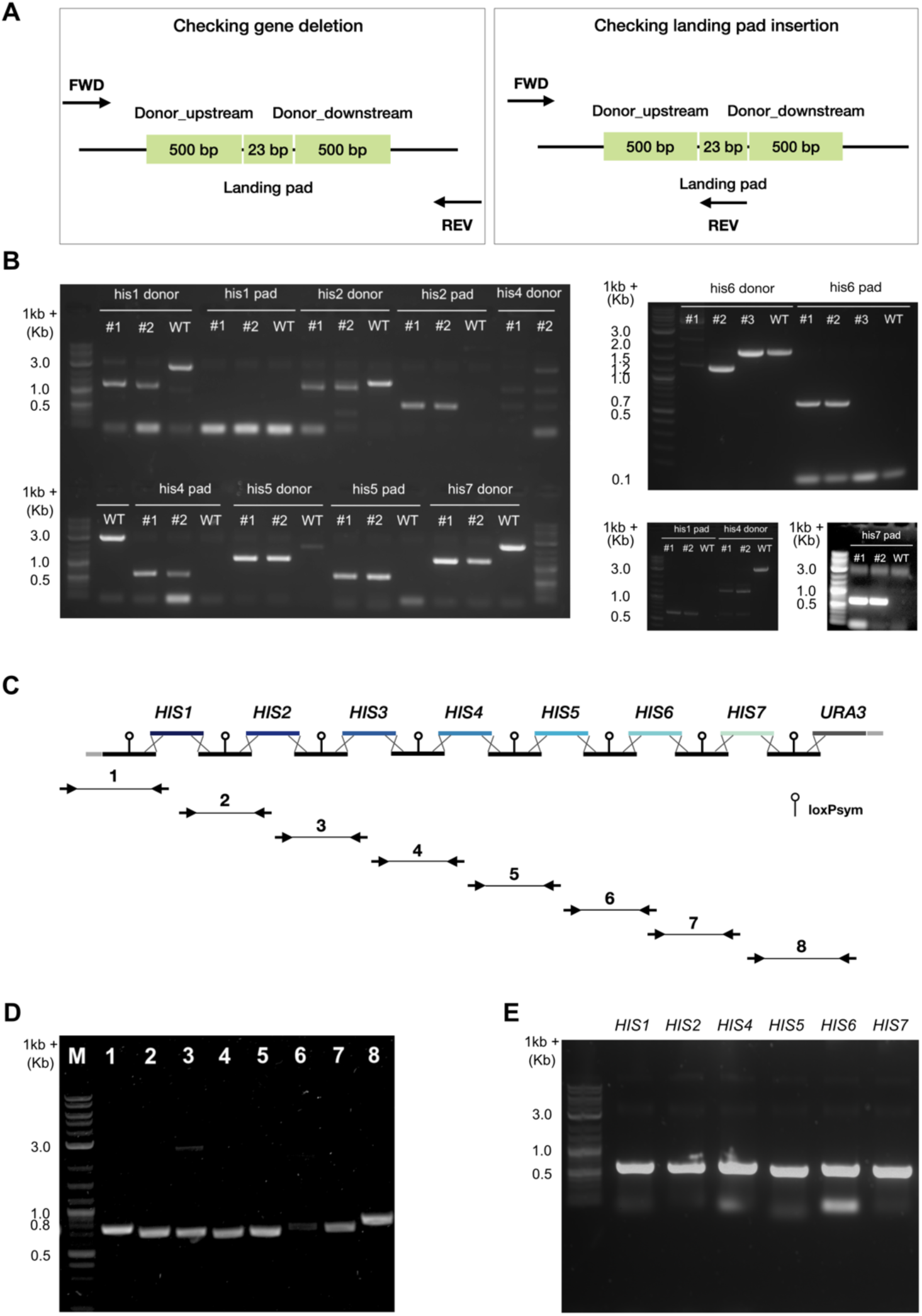
Junction PCR to verify gene deletions and synthetic module assembly. (A) Schematic of primer design to identify *HIS* gene deletion and landing pad insertion. (B) Colony PCR of transformants to identify *HIS* gene deletion. Numbers in white colour indicate the various tested colonies. (C) Schematic of synthetic *HIS* module assembly by yeast homology-dependent recombination and primer design to identify synthetic *HIS* module integration. Arrows in black represent the primers targeting at the junctions for PCR. Lines connected the arrows with numbers above indicate the various tested junctions. (D) Junction PCR from the genomic DNA of strain yXL052 for confirming the defragmented *HIS* cluster integration. (E) PCR from genomic DNA of strain yXL052 for confirming the landing pad insertion at the *HIS* gene deletion sites.

**Figure S3.**
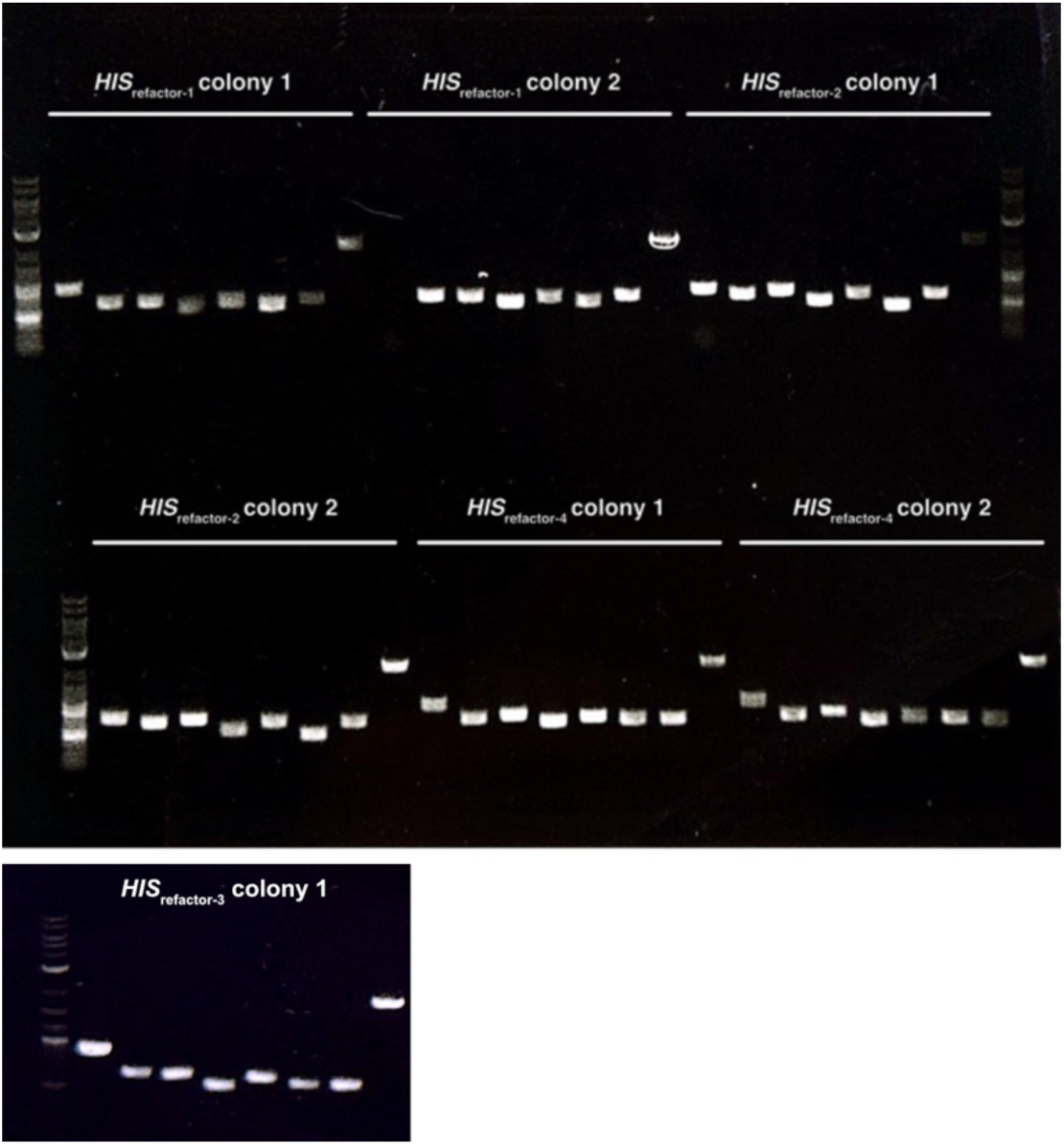
Junction PCR to verify the assembly of the refactored *HIS* modules. Transformants were randomly selected for checking the synthetic module assemblies. Eight junctions were checked for each strain.

**Figure S4.**
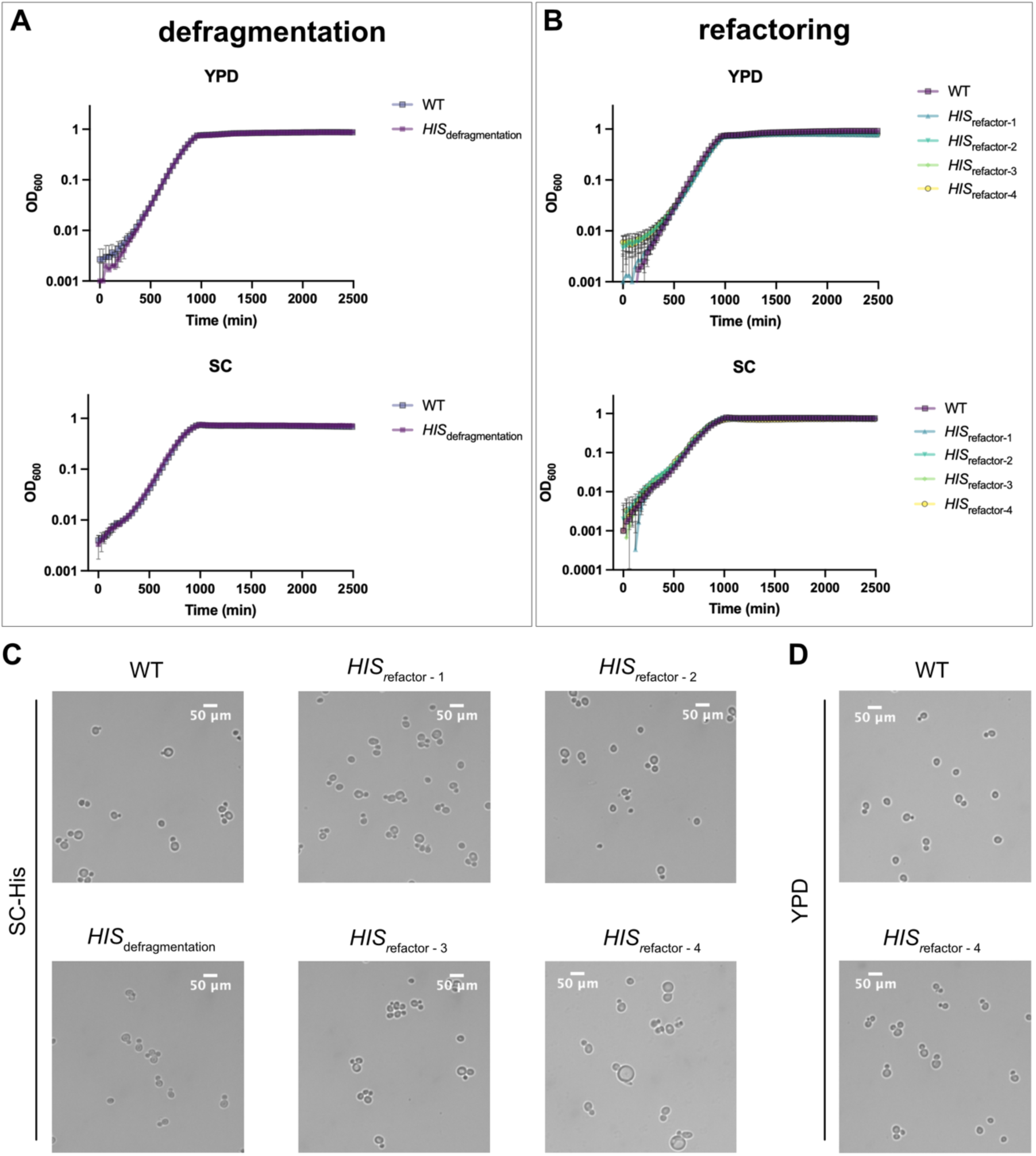
Physiological characterisation of the strains harbouring the defragmented and refactored *HIS* cluster. (A) Growth curves of the WT control strain (yXL014) and the strain harbouring the defragmented *HIS* module (yXL052) in YPD and SC media, n=4. (B) Growth curves of the WT control strain (yXL014) and the strains harbouring the refactored *HIS* modules (yXL214, yXL215, yXL269 and yXL219), in which *HIS* genes are driven by high (*HIS*refactor-1), medium (*HIS*refactor-2), low (*HIS*refactor-3) and mixed strength (*HIS*refactor-4) promoters respectively, in YPD and SC media, n=3. (C) Microscopy images of the WT control strain (yXL014) and strains harbouring the synthetic *HIS* modules from the overnight culture in SC-His. (D) Microscopy images of the WT control strain (yXL014) and the strain harbouring the refactored *HIS* cluster (*HIS*refactor-4) from the overnight culture in YPD.

**Figure S5.**
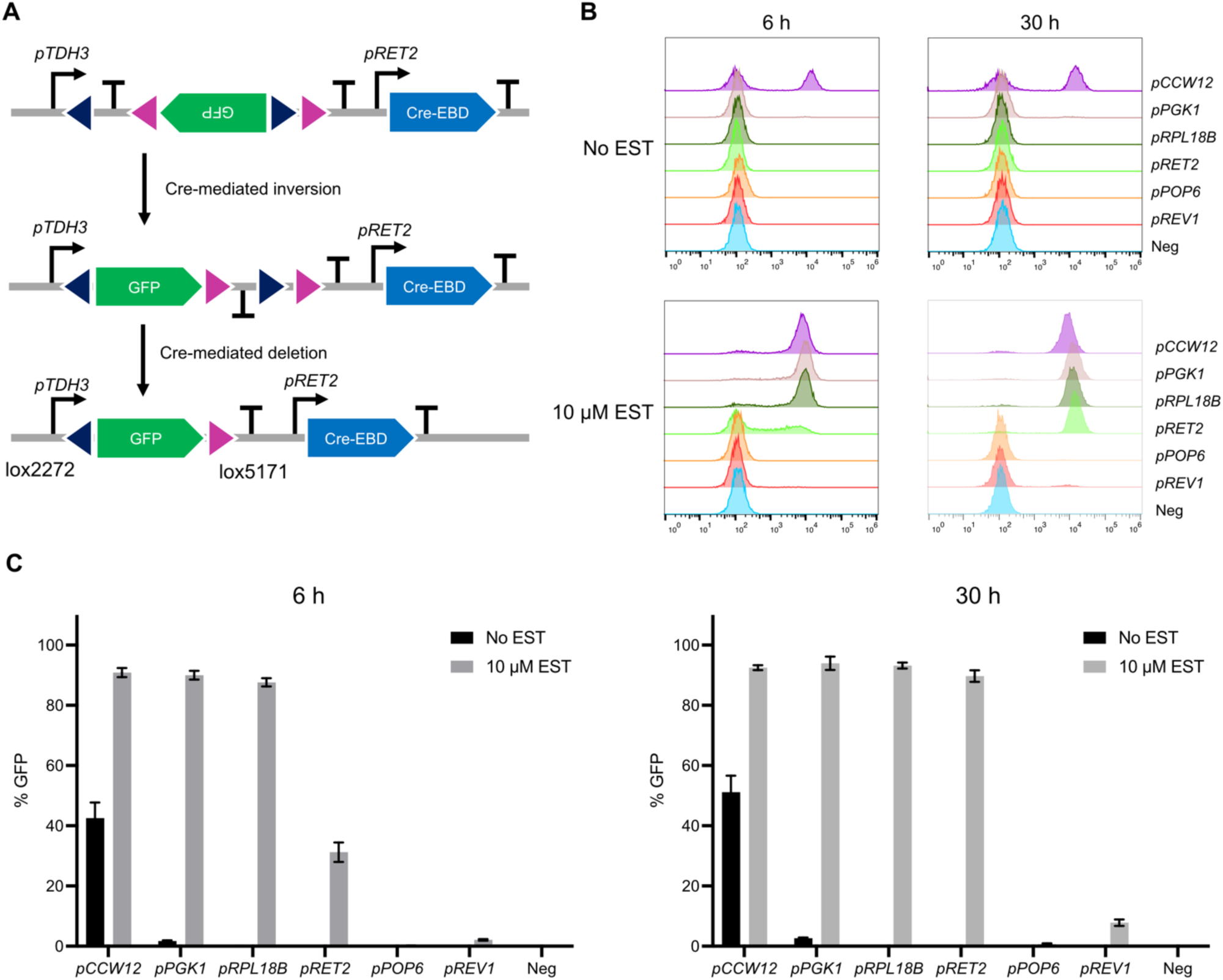
Characterisation of promoters for inducible Cre-EBD expression. (A) SCRaMbLE reporter constructs integrated at the *URA3* locus for characterisation of Cre-EBD expression promoters. RET2 promoter was finally selected as it showed zero leak and the cells fully convert to GFP^+^ over time. (B) Histograms of GFP fluorescence measured by flow cytometry after back dilution of saturated overnight cultures into fresh SC medium with and without 10 µM β-estradiol (EST) for 6 h (left) and 30 hours (right). (C) Percentage of the GFP^+^ yeast population after back dilution of saturated overnight cultures into fresh SC medium with and without 10 µM β-estradiol (EST) for 6 h (left) and 30 hours (right), n=3.

**Figure S6.**
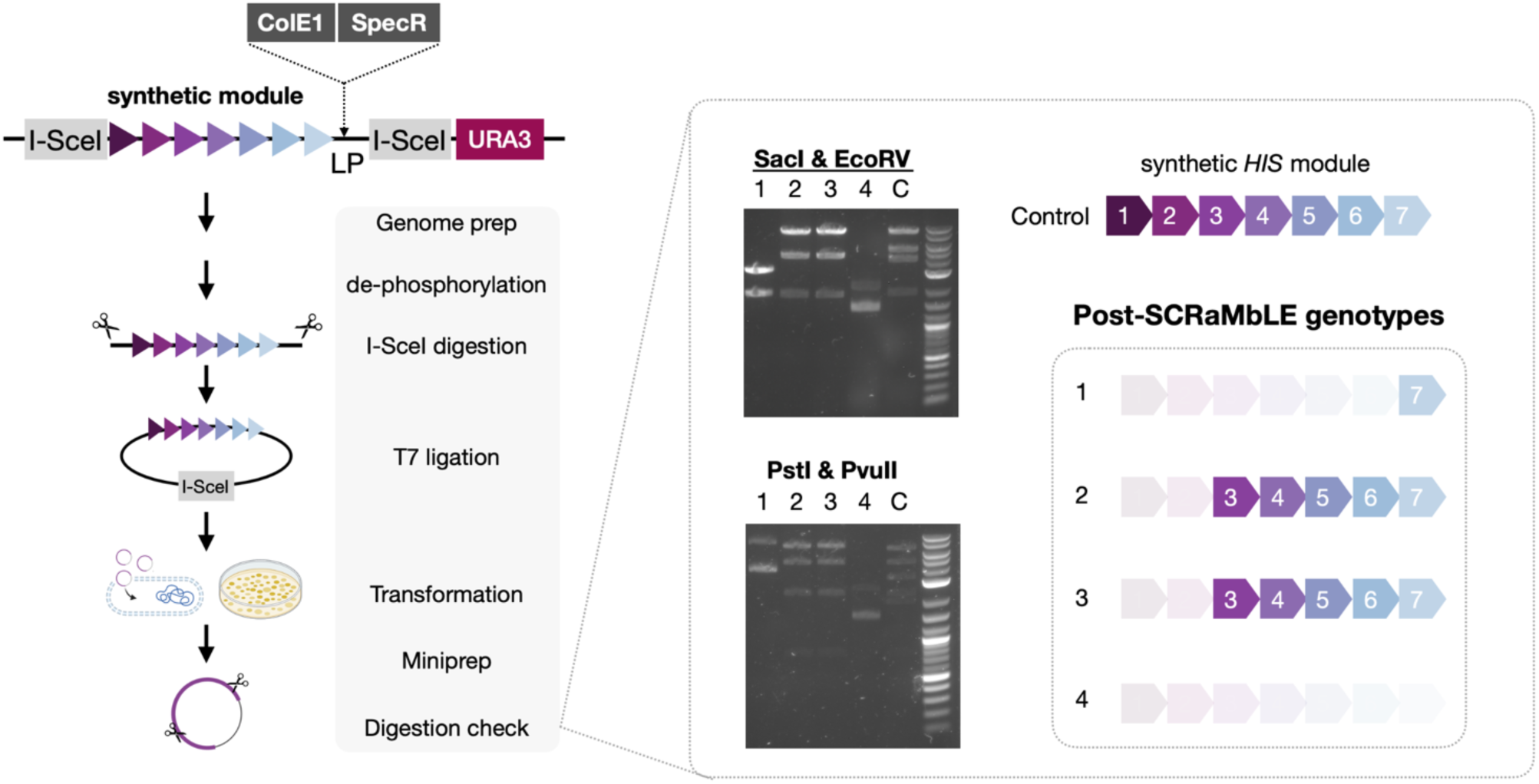
Analysis of the post-SCRaMbLE rearrangements using restriction digestion. Left: The ColE1 origin of replication and a selectable marker (*SpecR*) is inserted at the site of the landing pad (LP) between the synthetic module and the downstream I-SceI restriction site. The genomic region flanking by two I-SceI sites is then digested and ligated into a circular DNA through I-SceI digestion and T7 ligation. The ligation product is transformed into *E.coli* to enrich the circular plasmid containing the synthetic module. After miniprep, various combinations of restriction digestion are performed to identify the gene rearrangement patterns post SCRaMbLE. Right: Digestion results and schematic showing the corresponding genotypes from 4 post-SCRaMbLE strains and a control strain from the uninduced sample, labelling as “1, 2, 3, 4 and C” in black, respectively. Numbers in white represent 7 *HIS* genes (*HIS1* to *HIS7*). Arrows indicate the direction of gene transcription. Some icons for illustrating the workflow are generated with Biorender.com.

**Figure S7.**
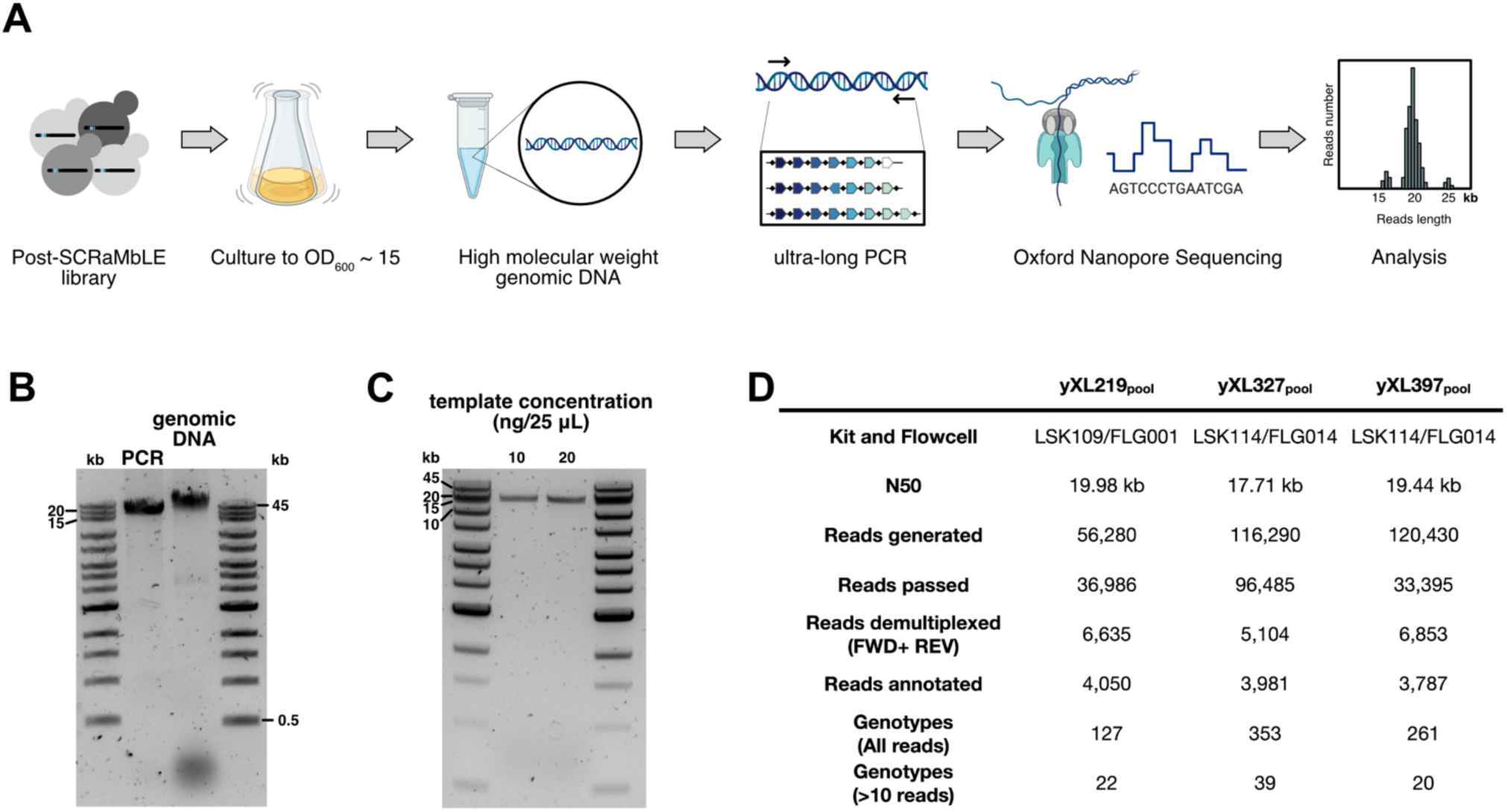
Workflow for POLAR-seq to identify gene rearrangements from the post-SCRaMbLE cell library. (A) The post-SCRaMbLE yeast cell library sorted through FACS are grown overnight to saturation (OD600 ∼15) in appropriate media in a baffled flask. 50 mL cell culture are collected to isolate whole genomic DNA with high molecular weight (HMW), which is next used as the template in long PCR to obtain amplicons of the rearranged synthetic module for nanopore sequencing. Long reads of the amplicons are detected by identifying the primer sequences and further annotated by a custom python script^26^. (B) Gel image showing the isolated HMW DNA of the post-SCRaMbLE library (yXL397pool) and amplification of the rearranged synthetic *HIS* module using this isolated HMW DNA as the template. (C) Optimisation of template concentration for ultra-long PCR. 6 μL of each PCR product was checked by electrophoresis. (D) Statistics of nanopore sequencing runs of sample yXL219pool, yXL327pool and yXL397pool. Some icons in panel A are created with BioRender.com.

**Figure S8.**
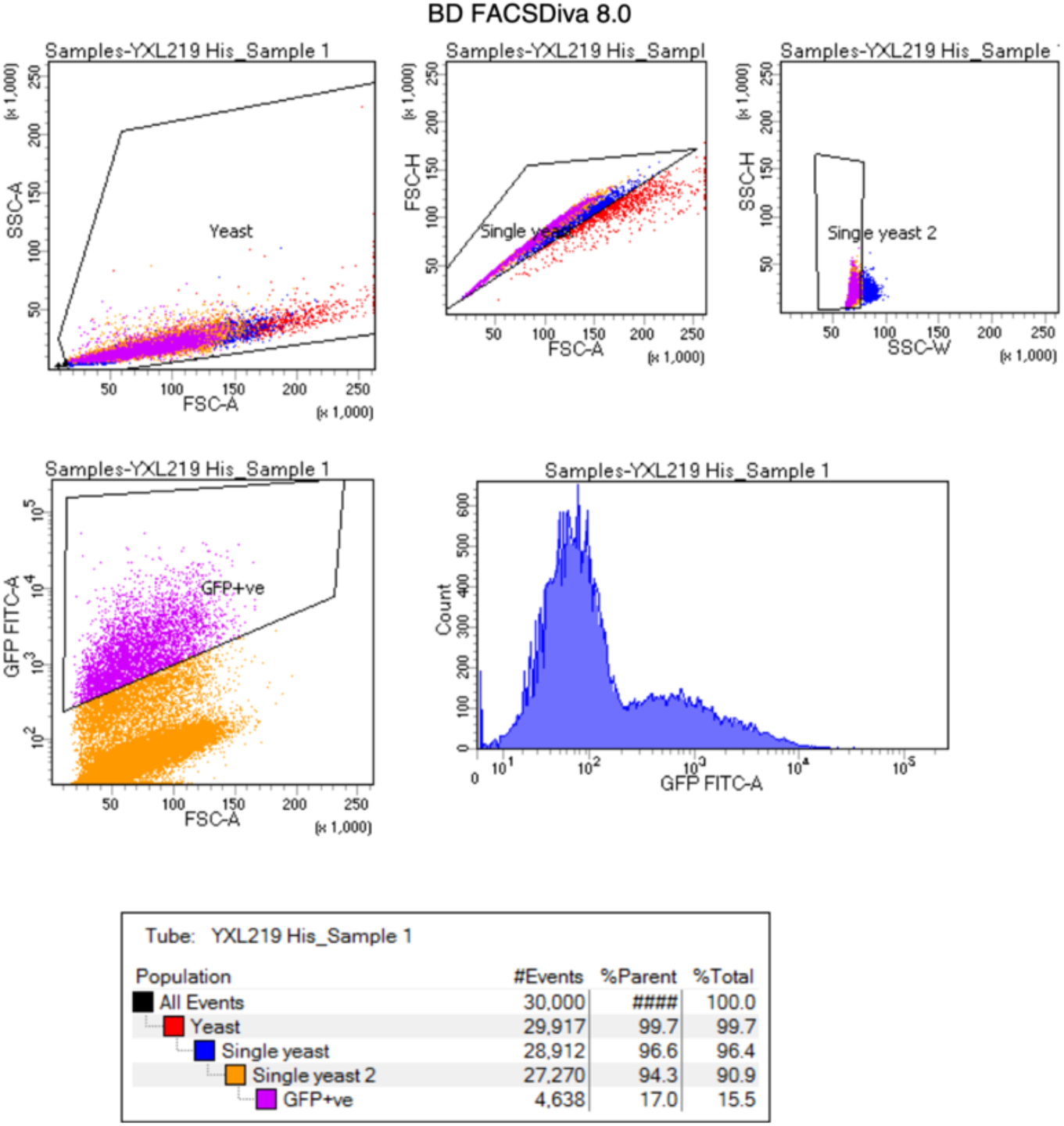
FACS sorting of post-SCRaMbLE cells using SCOUT. Yeast cells, single cells and GFP^+^ cells were gated manually. Percentage of these gated populations is shown in the table at the bottom.

**Figure S9.**
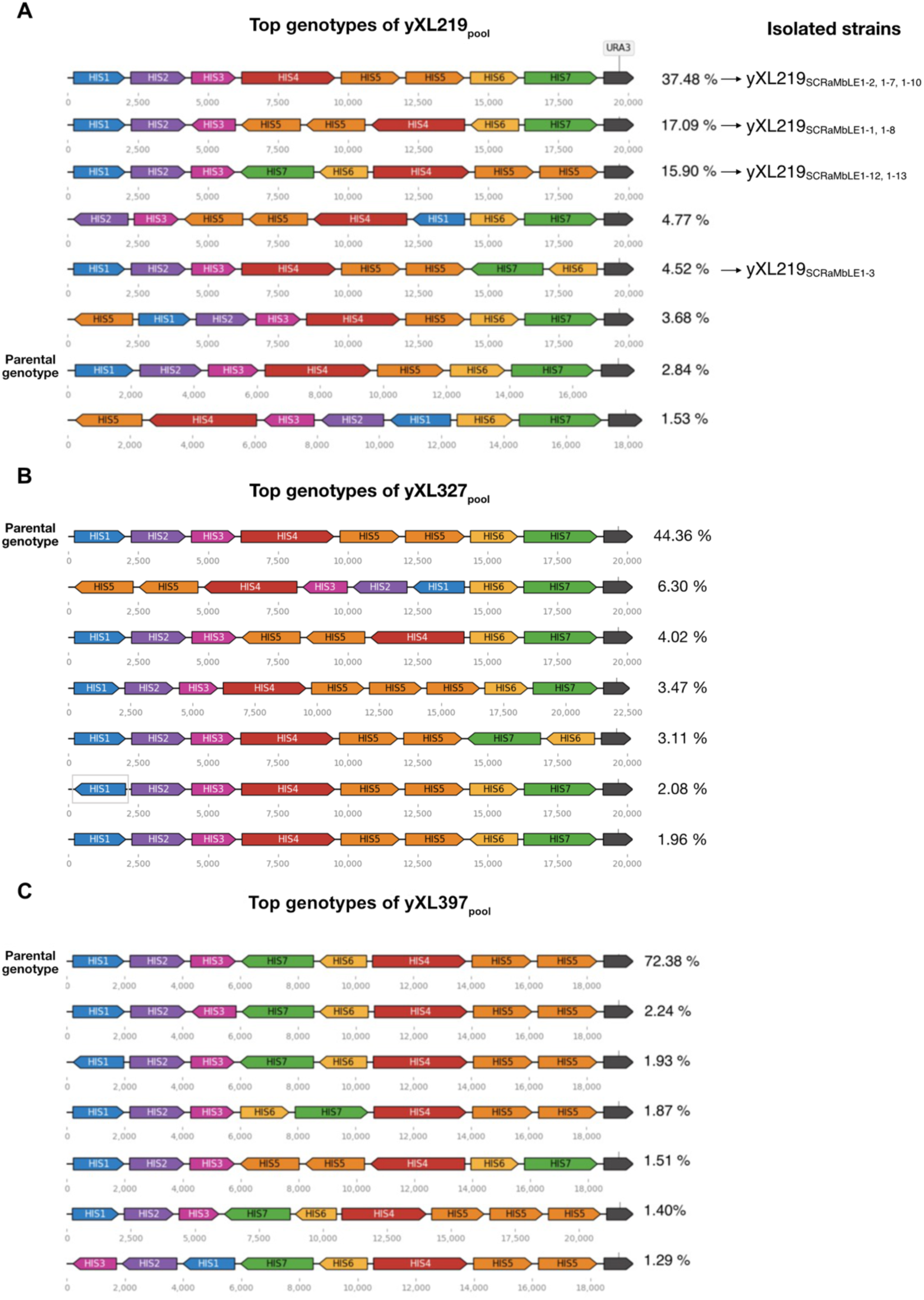
The most abundant genotypes identified from post-SCRaMbLE libraries (A) yXL219pool, (B) yXL327pool and (C) yXL397pool. Transcription units (TUs) of the genes are illustrated in different colours, with their transcription direction indicated by arrows. Reads frequency of each genotype is shown on the right. Numbers in grey below each genotype annotate the positions (in bp) of each TU within the synthetic module. Strains isolated from yXL219pool with their genotypes identified by nanopore sequencing are indicated by arrows on the right.

**Figure S10.**
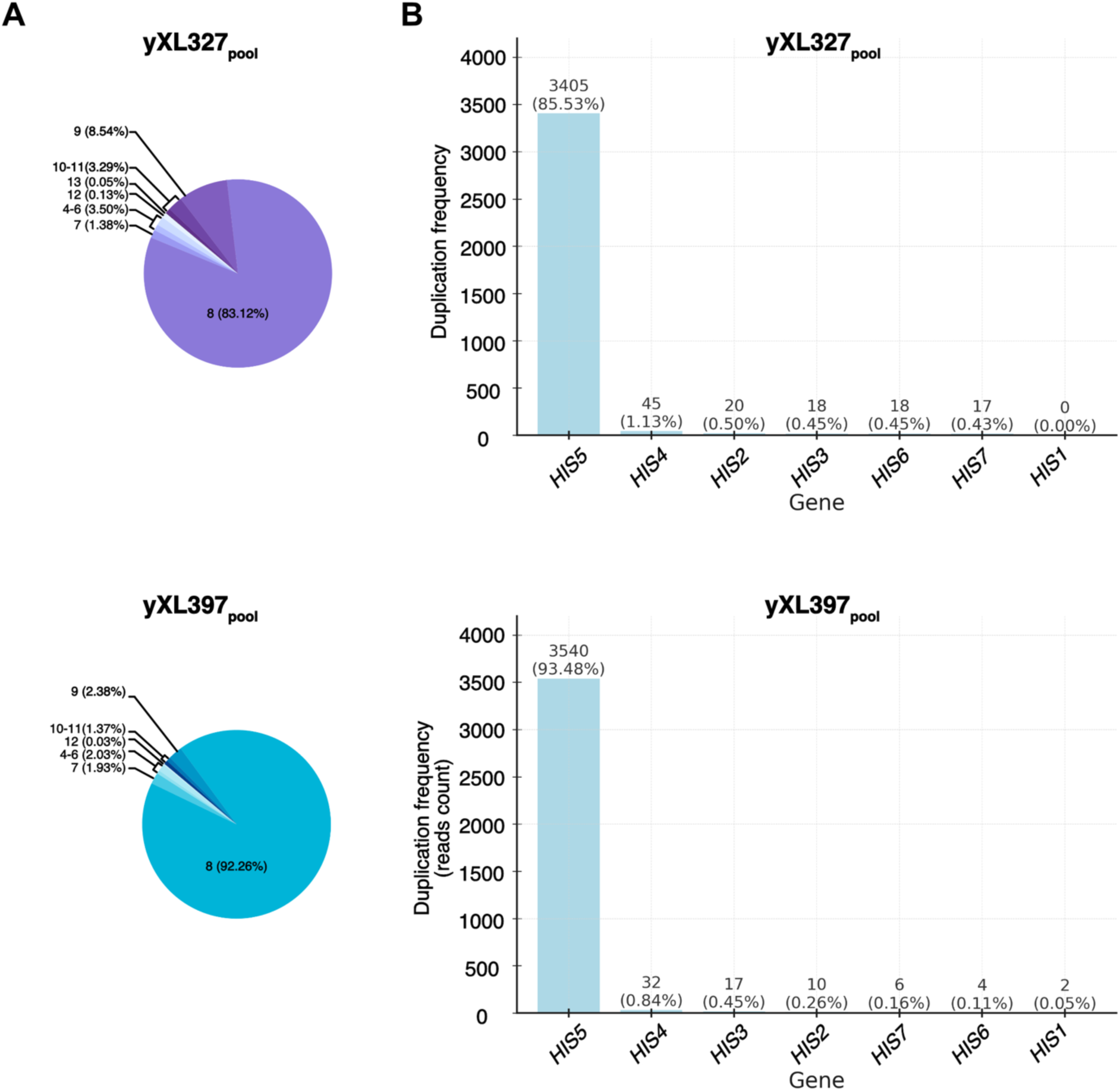
Genotypic composition of the post-SCRaMbLE libraries in the second round of SCRaMbLE. (A) Pie charts showing the number of genes identified from each distinct genotype and the percentage of total read count for each genotype featured with specific gene numbers. (B) Bar charts displaying the duplication frequency of each gene in the detected genotypes, calculated by the sum of reads count for genotypes detected with the duplicated genes dividing by the total read count in respective datasets.

## Supplementary Tables

**Supplementary Table 1.**
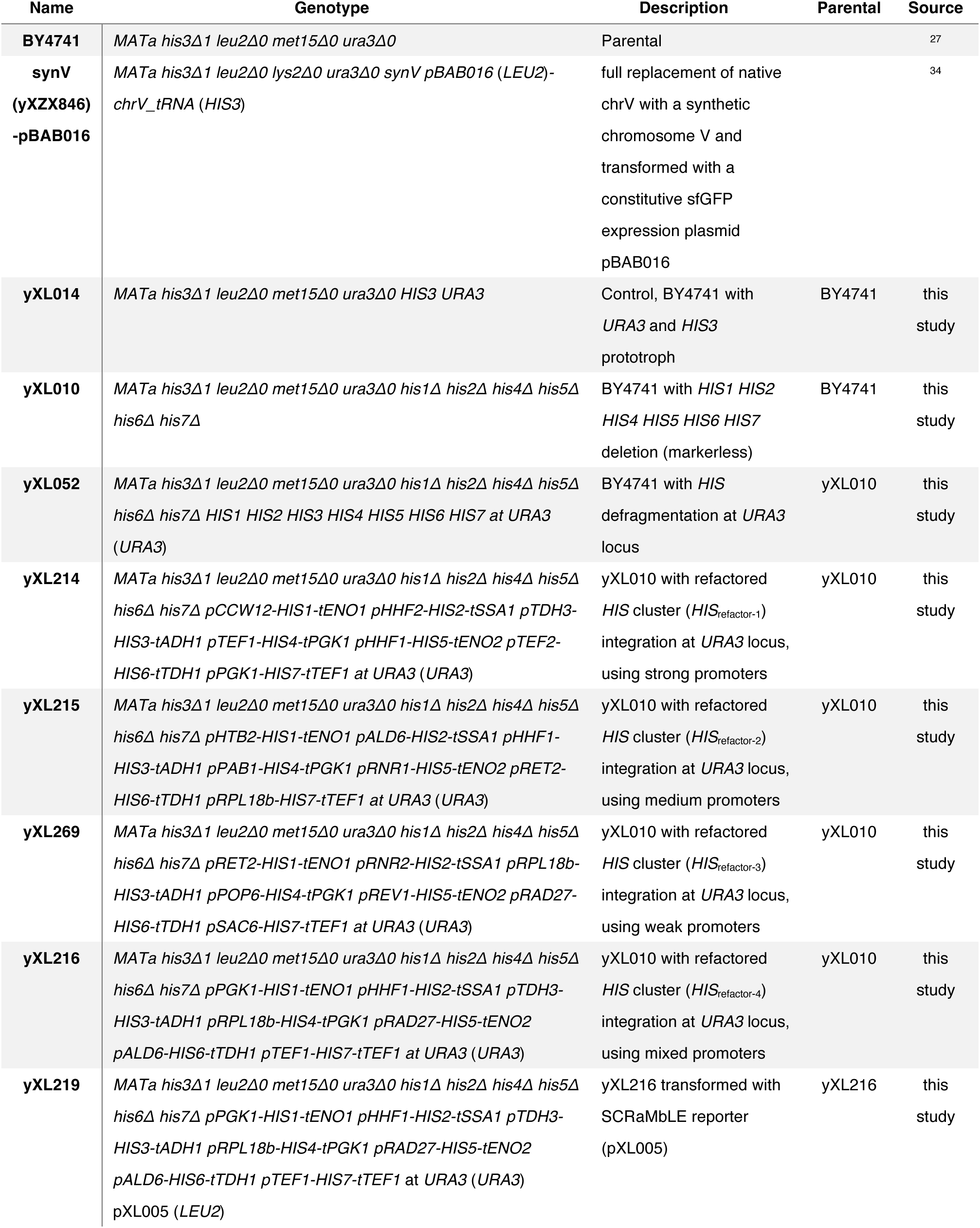
List of strains generated in this study.

**Supplementary Table 2.**
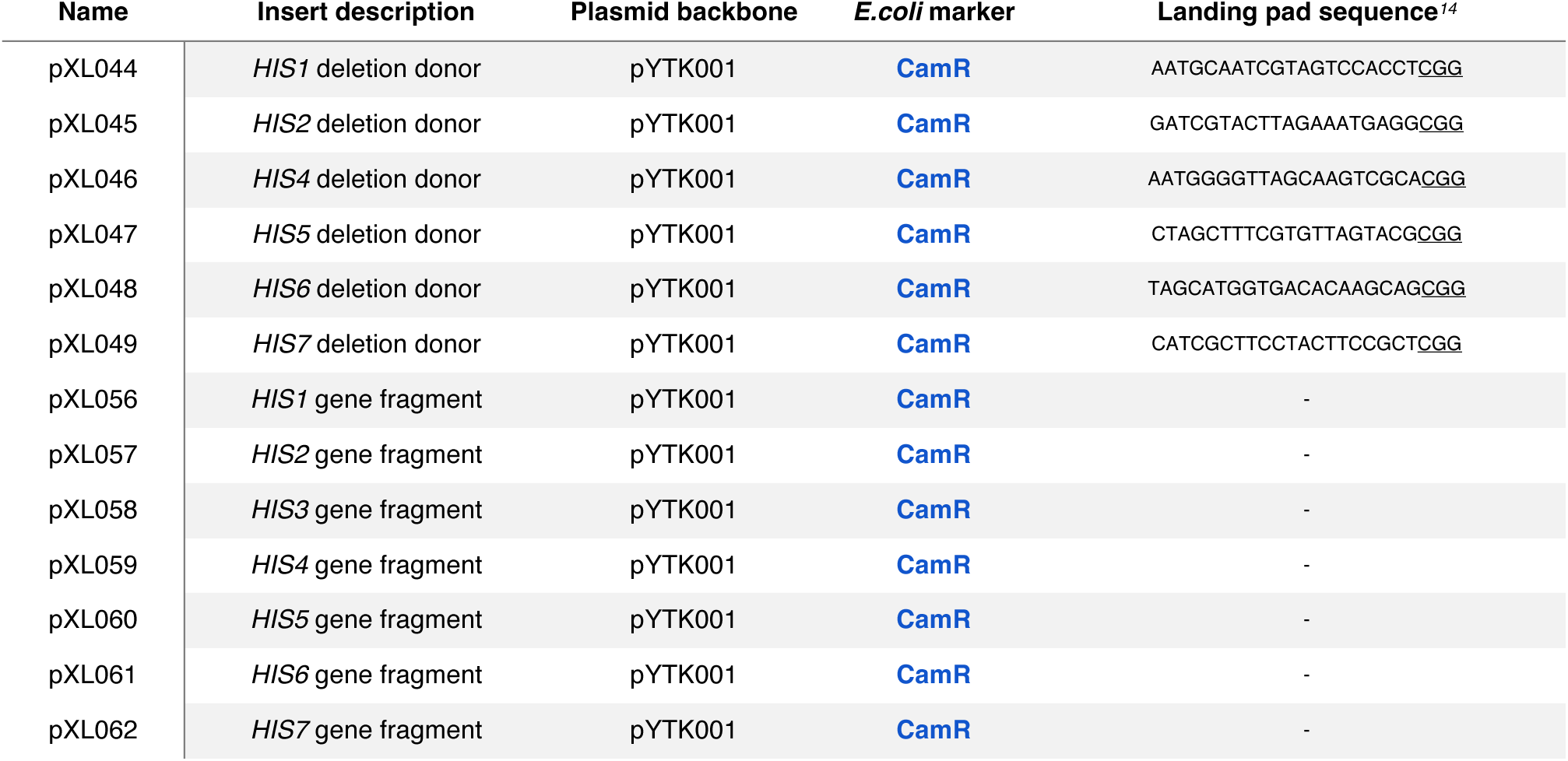
List of gap repair donor and gene fragment plasmids used for defragmentation of *HIS* biosynthesis in this study. To ease future engineering of the sites left behind by gene deletion, we substituted each deleted sequences with an individual 23 bp ‘landing pad’^14^ that encodes a unique CRISPR/Cas9 target sequence. Landing pad sequences are shown in the right column with protospacer adjacent motif (PAM) sequence highlighted by underlining.

**Supplementary Table 3.**
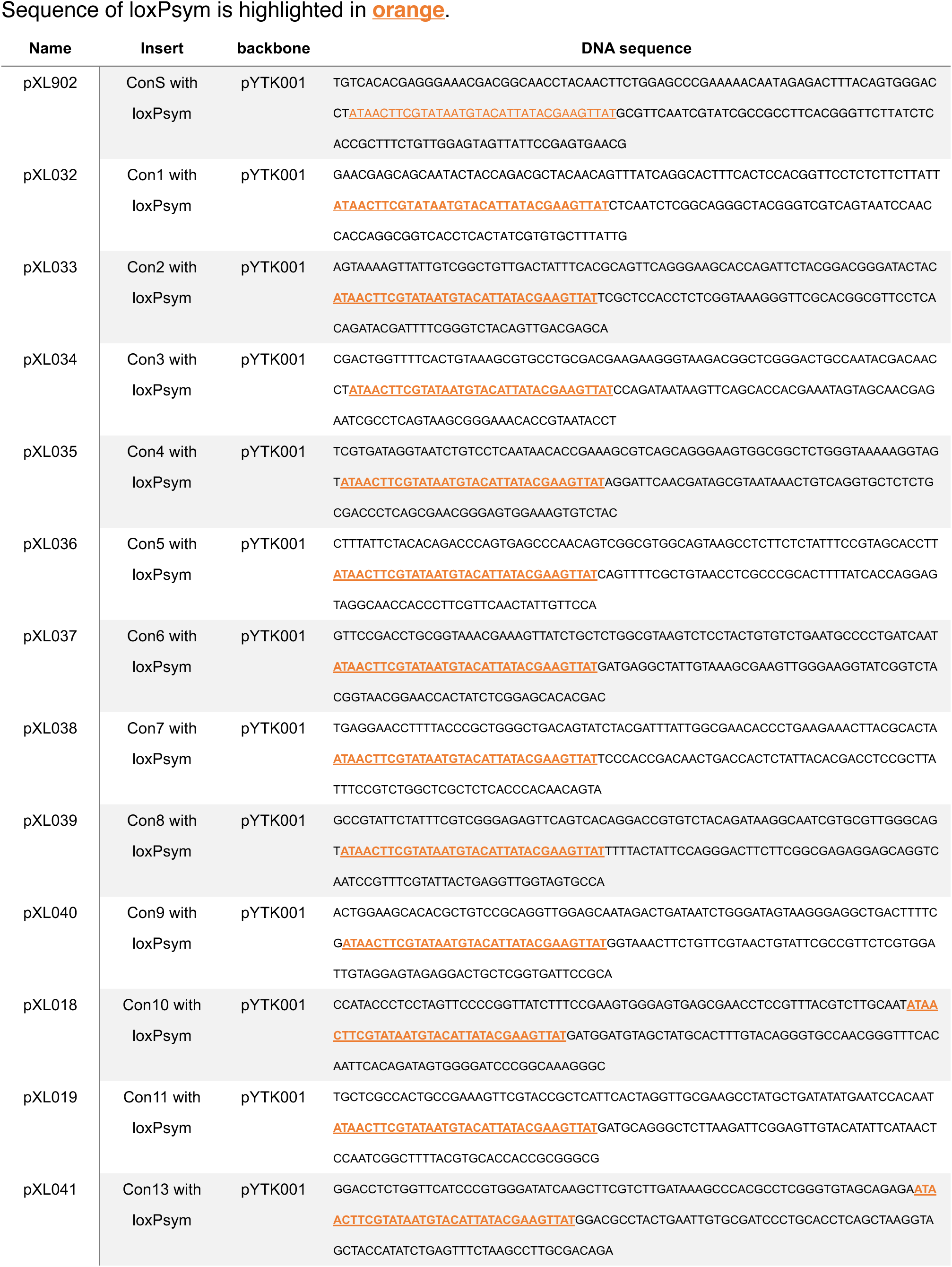
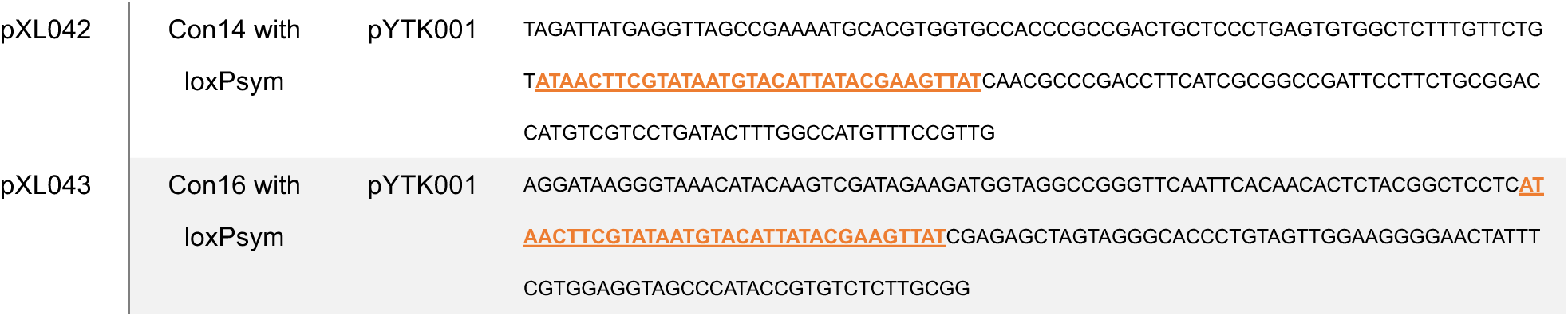
List of linker plasmids used in this study.

**Supplementary Table 4.**
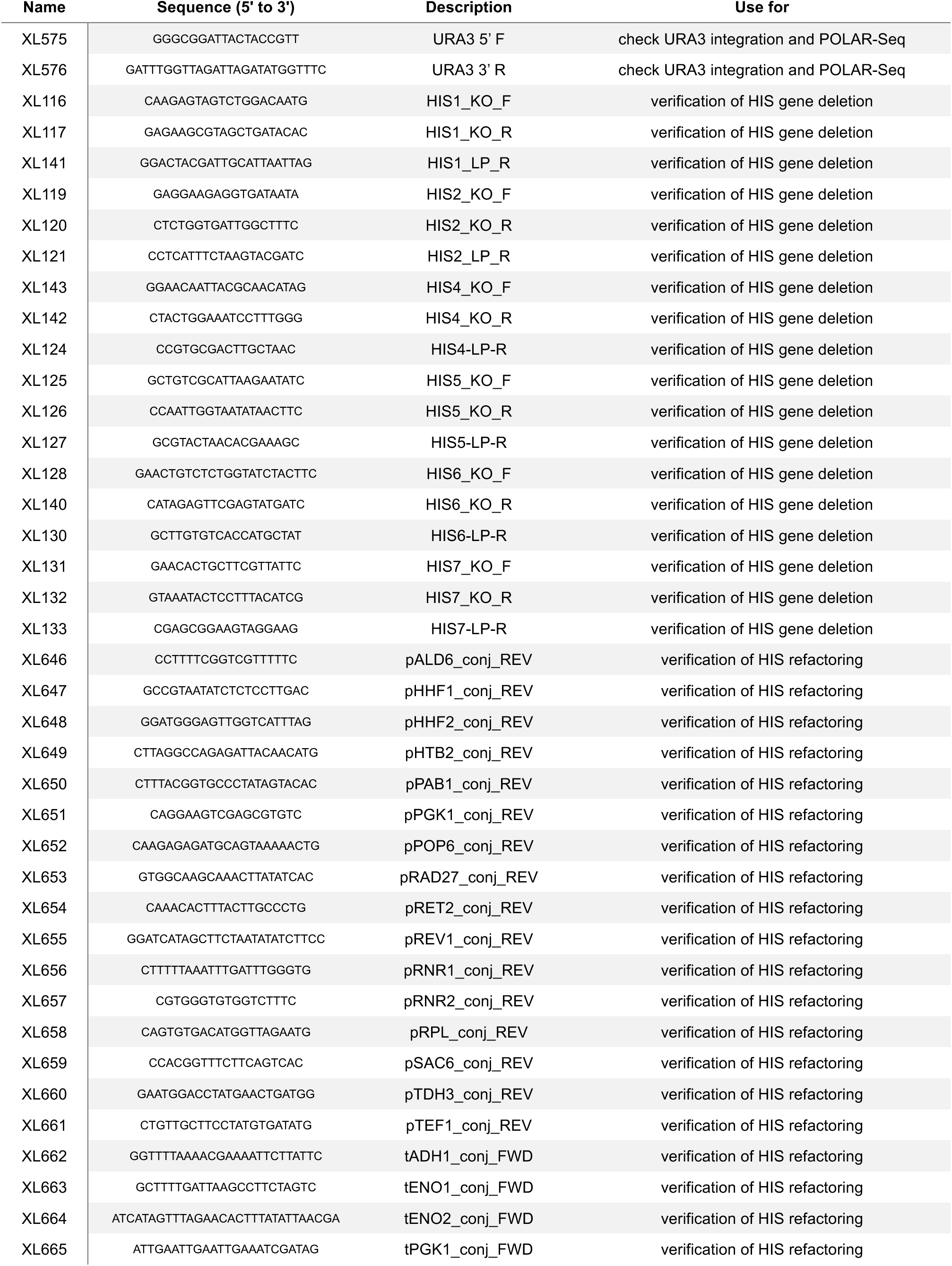
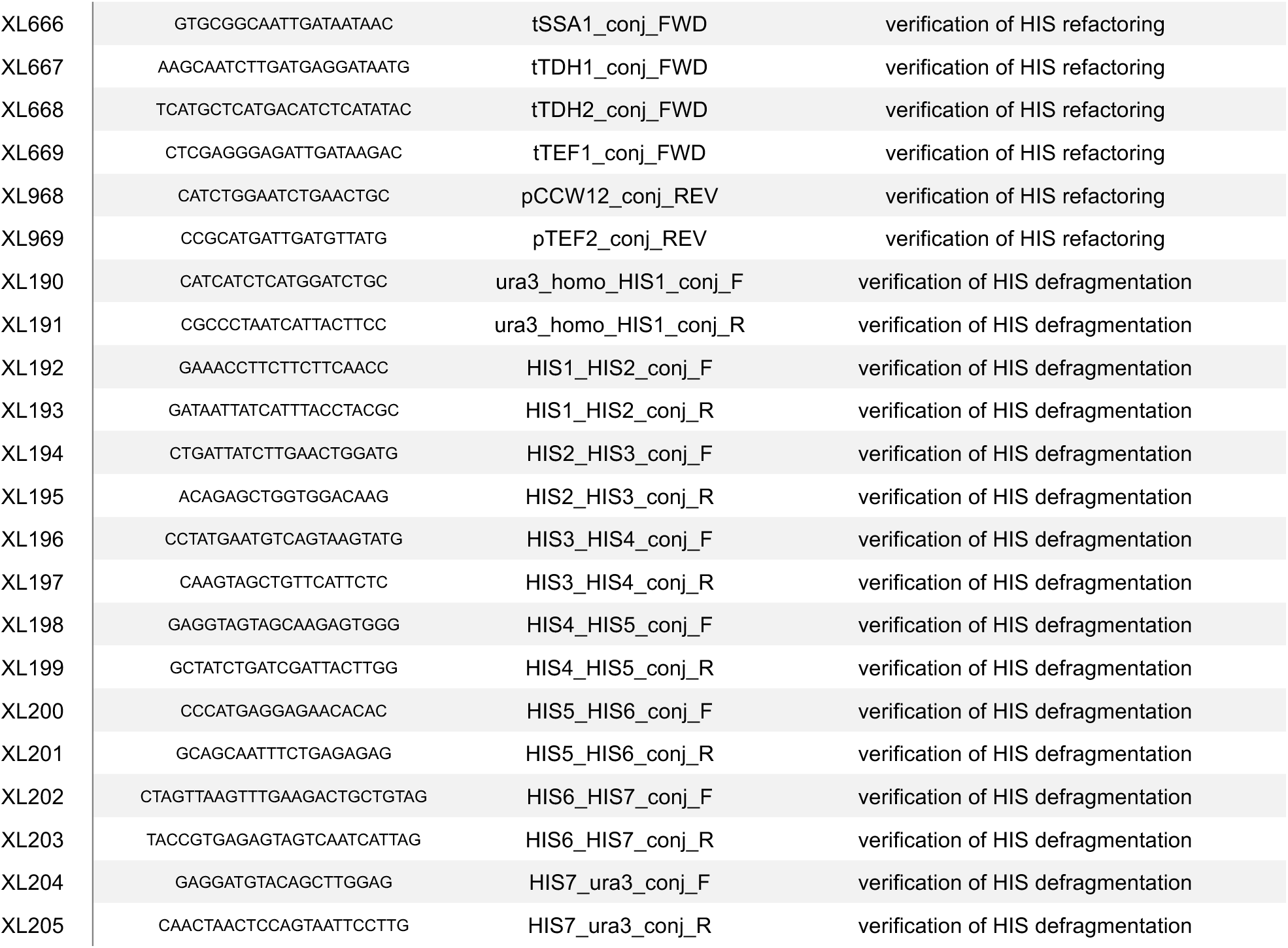
List of oligos used in this study.

**Supplementary Table 5.**
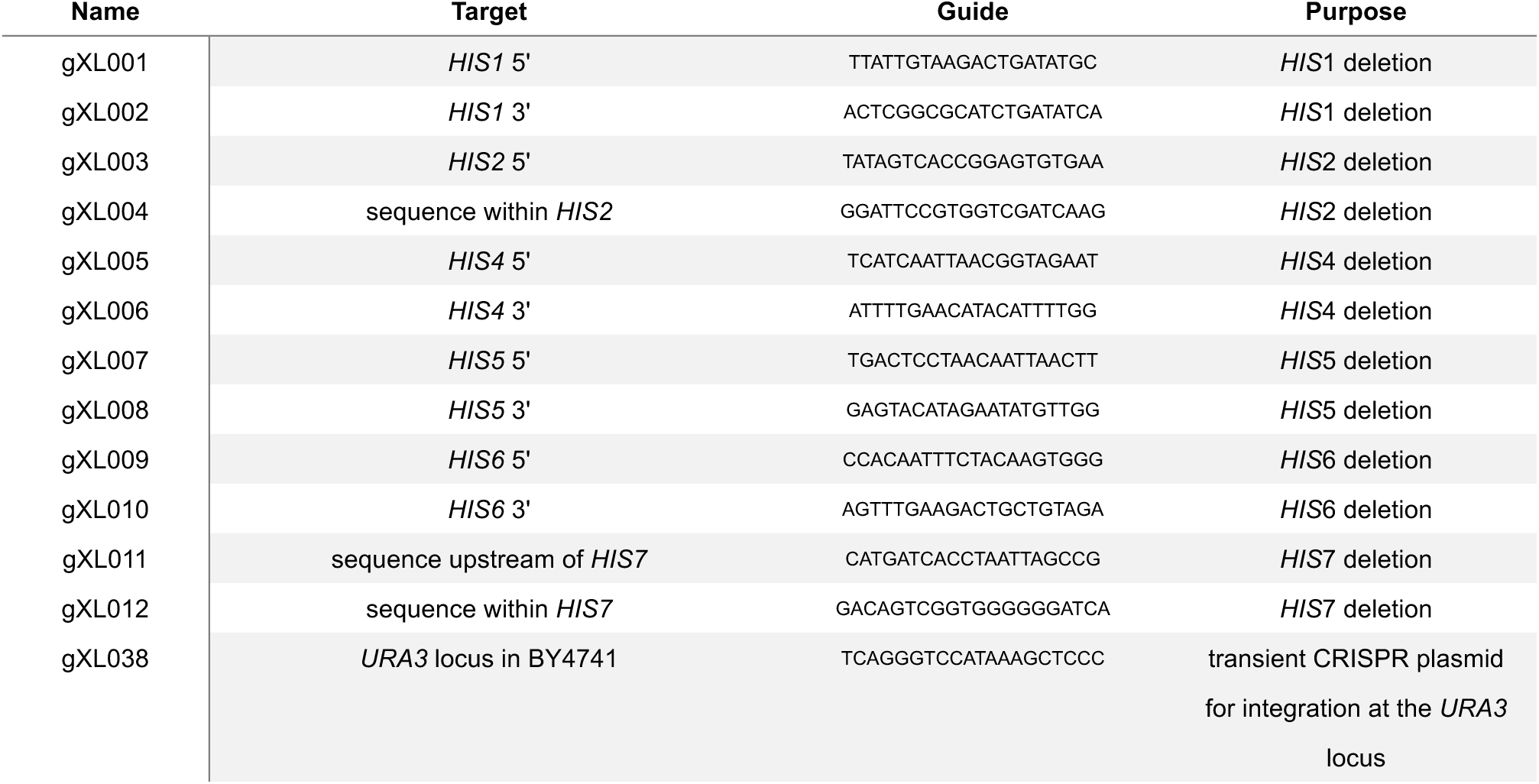
List of gRNAs used in this study.

**Supplementary Table 6.** List of parts in YTK format used in this study.

| Name | Part type | Part description | Source* | *E.coli* marker |
| --- | --- | --- | --- | --- |
| pXL009 | 3 | *HIS1* | PCR from BY4741 genome | CamR |
| pXL010 | 3 | *HIS2* | PCR from BY4741 genome | CamR |
| pWS942 | 3 | *HIS3* | Will Shaw | CamR |
| pXL011 | 3 | *HIS4_G599A* | PCR from BY4741 genome | CamR |
| pXL012 | 3 | *HIS5* | PCR from BY4741 genome | CamR |
| pXL013 | 3 | *HIS6* | PCR from BY4741 genome | CamR |
| pXL014 | 3 | *HIS7_A1341G* | PCR from BY4741 genome | CamR |
| pJCH021 | 2 | *pSCW11* | Jack Ho | CamR |
| pJCH022 | 3 | *cre* | Jack Ho | CamR |
| pTMP059 | 4 | *tTEF1* | YTK addition | CamR |

\* Tom Ellis Lab members who provided the parts are acknowledged by name.

**Supplementary Table 7.**
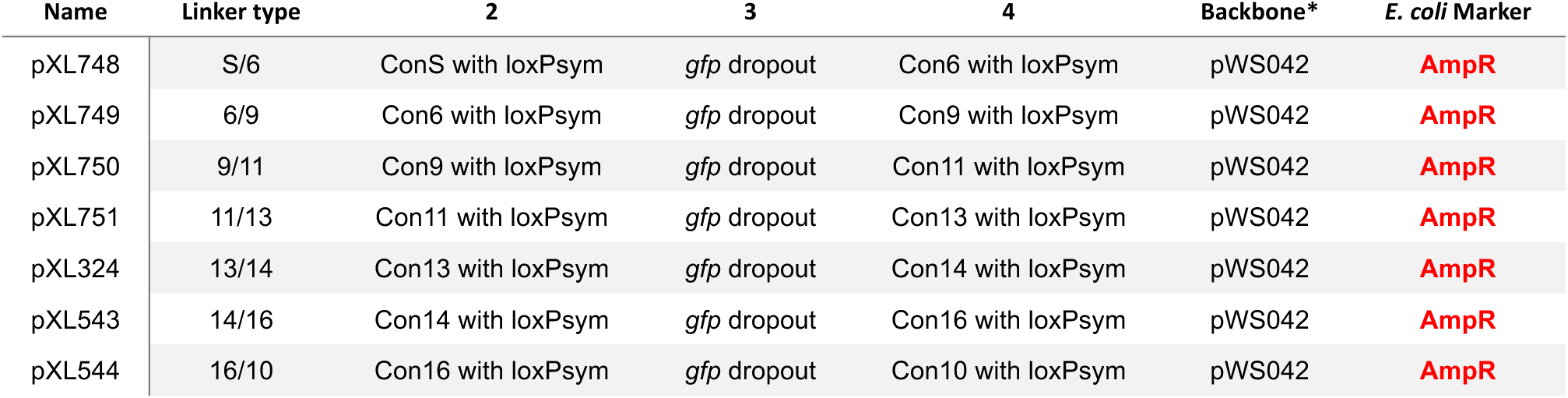
List of pre-assembled linker vectors used in this study.

| Name | Linker type | 2 | 3 | 4 | Backbone* | *E. coli* Marker |
| --- | --- | --- | --- | --- | --- | --- |
| pXL748 | S/6 | ConS with loxPsym | *gfp* dropout | Con6 with loxPsym | pWS042 | **AmpR** |
| pXL749 | 6/9 | Con6 with loxPsym | *gfp* dropout | Con9 with loxPsym | pWS042 | **AmpR** |
| pXL750 | 9/11 | Con9 with loxPsym | *gfp* dropout | Con11 with loxPsym | pWS042 | **AmpR** |
| pXL751 | 11/13 | Con11 with loxPsym | *gfp* dropout | Con13 with loxPsym | pWS042 | **AmpR** |
| pXL324 | 13/14 | Con13 with loxPsym | *gfp* dropout | Con14 with loxPsym | pWS042 | **AmpR** |
| pXL543 | 14/16 | Con14 with loxPsym | *gfp* dropout | Con16 with loxPsym | pWS042 | **AmpR** |
| pXL544 | 16/10 | Con16 with loxPsym | *gfp* dropout | Con10 with loxPsym | pWS042 | **AmpR** |

**Supplementary Table 8.** List of cassettes used in this study.

| Name | Cassette<br>type | 2 | 3a-1 | 3a-2 | 3b | 4 | Vector | *E. coli*<br>Marker | Yeast<br>Marker | Yeast<br>Vector |
| --- | --- | --- | --- | --- | --- | --- | --- | --- | --- | --- |
| pXL001 | 1/E | pJCH021 |  | pJCH022 |  | pYTK051 | pWS042 | **AmpR** |  |  |
| pXL003 | S/1 | pTDH3-lox2272-tCYC1-lox5171-mGFPmut2-lox2272-lox5171-tADH1 |  |  |  |  | pWS041 | **AmpR** |  |  |
| pXL445 | S/6 | pYTK016 |  | pXL009 |  | pYTK051 | pXL748 | **AmpR** |  |  |
| pXL446 | 6/9 | pYTK018 |  | pXL010 |  | pYTK052 | pXL749 | **AmpR** |  |  |
| pXL447 | 9/11 | pYTK015 |  | pWS942 |  | pYTK053 | pXL750 | **AmpR** |  |  |
| pXL448 | 11/13 | pYTK019 |  | pXL011 |  | pYTK054 | pXL751 | **AmpR** |  |  |
| pXL449 | 13/14 | pYTK021 |  | pXL012 |  | pYTK055 | pXL324 | **AmpR** |  |  |
| pXL558 | 14/16 | pYTK020 |  | pXL013 |  | pYTK056 | pXL543 | **AmpR** |  |  |
| pXL562 | 16/10 | pYTK017 |  | pXL014 |  | pTMP059 | pXL544 | **AmpR** |  |  |
| pXL450 | S/6 | pYTK010 |  | pXL009 |  | pYTK051 | pXL748 | **AmpR** |  |  |
| pXL451 | 6/9 | pYTK012 |  | pXL010 |  | pYTK052 | pXL749 | **AmpR** |  |  |
| pXL452 | 9/11 | pYTK009 |  | pWS942 |  | pYTK053 | pXL750 | **AmpR** |  |  |
| pXL453 | 11/13 | pYTK013 |  | pXL011 |  | pYTK054 | pXL751 | **AmpR** |  |  |
| pXL454 | 13/14 | pYTK015 |  | pXL012 |  | pYTK055 | pXL324 | **AmpR** |  |  |
| pXL559 | 14/16 | pYTK014 |  | pXL013 |  | pYTK056 | pXL543 | **AmpR** |  |  |
| pXL563 | 16/10 | pYTK011 |  | pXL014 |  | pTMP059 | pXL544 | **AmpR** |  |  |
| pXL455 | S/6 | pYTK020 |  | pXL009 |  | pYTK051 | pXL748 | **AmpR** |  |  |
| pXL456 | 6/9 | pYTK023 |  | pXL010 |  | pYTK052 | pXL749 | **AmpR** |  |  |
| pXL457 | 9/11 | pYTK017 |  | pWS942 |  | pYTK053 | pXL750 | **AmpR** |  |  |
| pXL458 | 11/13 | pYTK024 |  | pXL011 |  | pYTK054 | pXL751 | **AmpR** |  |  |
| pXL459 | 13/14 | pYTK027 |  | pXL012 |  | pYTK055 | pXL324 | **AmpR** |  |  |
| pXL560 | 14/16 | pYTK025 |  | pXL013 |  | pYTK056 | pXL543 | **AmpR** |  |  |
| pXL564 | 16/10 | pYTK022 |  | pXL014 |  | pTMP059 | pXL544 | **AmpR** |  |  |
| pXL460 | S/6 | pYTK011 |  | pXL009 |  | pYTK051 | pXL748 | **AmpR** |  |  |
| pXL461 | 6/9 | pYTK015 |  | pXL010 |  | pYTK052 | pXL749 | **AmpR** |  |  |
| pXL462 | 9/11 | pYTK009 |  | pWS942 |  | pYTK053 | pXL750 | **AmpR** |  |  |
| pXL463 | 11/13 | pYTK017 |  | pXL011 |  | pYTK054 | pXL751 | **AmpR** |  |  |
| pXL464 | 13/14 | pYTK025 |  | pXL012 |  | pYTK055 | pXL324 | **AmpR** |  |  |
| pXL561 | 14/16 | pYTK018 |  | pXL013 |  | pYTK056 | pXL543 | **AmpR** |  |  |
| pXL565 | 16/10 | pYTK013 |  | pXL014 |  | pTMP059 | pXL544 | **AmpR** |  |  |

**Supplementary Table 9.** List of multigene cassettes used in this study.

| Name | Cassette type | 1 (vector) | 2 | 3 | 4 | 5 | 6 | *E. coli* Marker | Yeast Marker | Yeast Vector |
| --- | --- | --- | --- | --- | --- | --- | --- | --- | --- | --- |
| pXL004 | o | pWS036 | pXL003 | pXL001 |  |  |  | KanR | LEU2 | low-copy |
| pXL005 | o | pWS036 | pXL003 | pWS2409 |  |  |  | KanR | LEU2 | low-copy |

**Supplementary Table 10.** Statistics on multiplexed nanopore sequencing (R1-5).

| Raw reads |  |  |  |  |  |
| --- | --- | --- | --- | --- | --- |
|  | Reads | Throughput (Mbp) | Mean read length (kb) | N50 (kb) | Genome coverage |
| **R1** | 121,607 | 1,208 | 9.9 | 20.8 | 99.8 |
| **R2** | 121,791 | 1,133 | 9.3 | 18.6 | 93.6 |
| **R3** | 86,293 | 1,042 | 12.1 | 25.4 | 86.1 |
| **R4** | 131,687 | 1,370 | 10.4 | 21.0 | 113.2 |
| **R5** | 99,960 | 1,315 | 13.2 | 27.9 | 108.7 |

| Canu-corrected reads |  |  |  |  |  |
| --- | --- | --- | --- | --- | --- |
|  | Reads | Throughput (Mbp) | Mean read length (kb) | N50 (kb) | Genome coverage |
| **R1** | 11,821 | 486 | 41.1 | 45.5 | 40.2 |
| **R2** | 12,976 | 485 | 37.4 | 41.0 | 40.1 |
| **R3** | 11,263 | 485 | 43.0 | 46.7 | 40.1 |
| **R4** | 11,770 | 490 | 41.6 | 48.1 | 40.5 |
| **R5** | 10,728 | 492 | 45.9 | 53.8 | 40.7 |

